# Translational regulation of non-autonomous mitochondrial stress response promotes longevity

**DOI:** 10.1101/533695

**Authors:** Jianfeng Lan, Jarod Rollins, Di Wu, Xiao Zang, Lina Zou, Zi Wang, Chang Ye, Zixing Wu, Pankaj Kapahi, Aric N. Rogers, Di Chen

## Abstract

Inhibition of mRNA translation delays aging, but the underlying mechanisms remain underexplored. Mutations in both DAF-2 (IGF-1 receptor) and RSKS-1 (ribosomal S6 kinase/S6K) cause synergistic lifespan extension in *C. elegans*. To understand the roles of S6K-mediated translational regulation in this process, we performed genome-wide translational profiling and genetic screens to identify genes that are not only regulated at the translational level in the *daf-2 rsks-1* mutant, but also affect lifespan. Inhibition of CYC-2.1 (cytochrome c) in the germline significantly extends lifespan through non-autonomous activation of the mitochondrial unfolded protein response (UPR^mt^) and AMP-activated kinase (AMPK) in the metabolic tissue. Furthermore, the RNA-binding protein GLD-1-mediated translational repression of cytochrome c in the germline is important for the non-autonomous activation of UPR^mt^ and synergistic longevity of the *daf-2 rsks-1* mutant. Together, these results illustrate a translationally regulated non-autonomous mitochondrial stress response mechanism in the modulation of lifespan by insulin-like signaling and S6K.

**Highlights:** - Longevity of the *daf-2 rsks-1* mutant is mediated by translational repression of ribosomal proteins and CYC-2.1/cytochrome c.
- Germline inhibition of *cyc-2.1* non-autonomously activates UPR^mt^ and AMPK to extend lifespan.
- GLD-1 represses germline *cyc-2.1* translation in the *daf-2 rsks-1* mutant.
- Translational regulation of *cyc-2.1* and UPR^mt^ contribute to the synergistic longevity of the *daf-2 rsks-1* mutant.

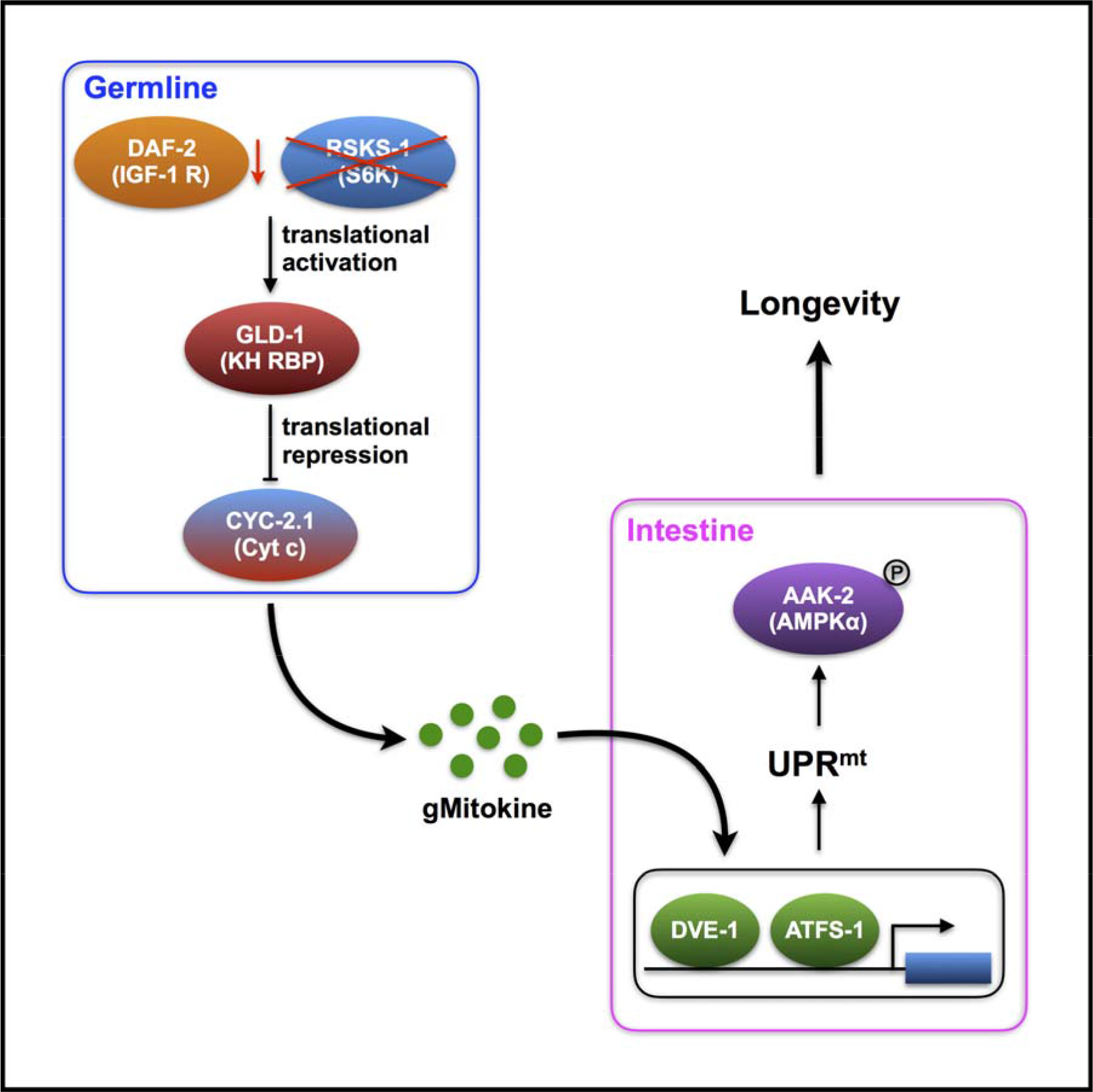

## Introduction

Aging can be genetically modulated by perturbation of insulin/insulin-like signaling (IIS), target of rapamycin (TOR) pathway and mitochondrial functions (Fontana et al., 2010; Kenyon, 2010; López-Otín et al., 2013). These genetic manipulations often lead to significant changes in gene expression both at transcriptional and translational levels. Inhibition of DAF-2, the *C. elegans* ortholog of the IGF-1 receptor, doubles adult lifespan via activating the DAF-16 (FOXO) transcriptional factor to regulate downstream genes involved in stress resistance, detoxification and metabolism (Kenyon et al., 1993; Kimura et al., 1997; Lin et al., 1997; McElwee, 2004; Murphy et al., 2003; Ogg et al., 1997). On the other hand, proteomics and translational state analysis revealed that the *daf-2* mutant shows alteration in translational states of gene expression extended survival (Depuydt et al., 2013; Dong et al., 2007; McColl et al., 2010; Stout et al., 2013).

Inhibition of the TOR pathway also significantly extends lifespan in many species (Kapahi et al., 2010). One important function of TOR is to regulate gene expression at the mRNA translation level through the ribosomal S6 kinase (S6K) and translational initiation factor 4E-binding protein (4E-BP), both of which have been shown to play important roles in aging (Hansen et al., 2007; Kapahi et al., 2004; Pan et al., 2007; Zid et al., 2009). Deletion mutants of *rsks-1*, which encodes the *C. elegans* ribosomal S6 kinase (S6K) ortholog, lead to significant changes in development, metabolism, reproduction and longevity (Hansen et al., 2007; Korta et al., 2012; Pan et al., 2007; Shi et al., 2013). Previous studies have identified multiple mediators of the *rsks-1* mutant, inhibition of which completely suppresses the longevity phenotype caused by *rsks-1* knockout mutants (Chen et al., 2009; McQuary et al., 2016; Selman et al., 2009; Seo et al., 2013; Sheaffer et al., 2008). However, genes translationally regulated by RSKS-1 that influence lifespan and the underlying molecular mechanisms remain to be characterized.

Serving as the key organelle in energy homeostasis, mitochondria play important but complex roles in aging. Mitochondrial dysregulation has been regarded as one of the major hallmarks of aging. On the other hand, mild perturbation of the mitochondrial electron transport chain (ETC) leads to significant lifespan extension in many species (Dillin et al., 2002; Houtkooper et al., 2014; Lee et al., 2002; López-Otín et al., 2013). Inhibition of mitochondrial ETC genes triggers the mitochondrial unfolded protein response (UPR^mt^) via transcriptional regulators such as DVE-1, UBL-5 and ATFS-1 (Haynes et al., 2007; Nargund et al., 2012). Intriguingly, perturbation of mitochondrial ETC functions in neurons releases a pro-longevity cue named mitokine to induce UPR^mt^ in the intestine, a distal metabolic tissue in worms, and ensures lifespan extension (Durieux et al., 2011). Further studies identified the neurotransmitter serotonin, neuropeptide FLP-2 and retromer-dependent Wnt signaling as the endocrine mediators of neurons to intestine non-autonomous mitochondrial stress response (Berendzen et al., 2016; Shao et al., 2016; Zhang et al., 2018). The trans-tissue mitochondrial stress response requires epigenetic modifications that ensure selective gene expression and prolonged longevity, and the epigenetic regulatory mechanisms are conserved in mammals (Merkwirth et al., 2016; Tian et al., 2016).

In order to study how insulin-like signaling interacts with the TOR pathway to modulate aging, we previously constructed a *daf-2 rsks-1* double mutant and observed a synergistic longevity phenotype. Further analysis demonstrated an AMPK-mediated positive feedback regulation of the DAF-16 transcriptional factor mechanism in the *daf-2 rsks-1* mutant (Chen et al., 2013). However, genes that are translationally regulated in the *daf-2 rsks-1* mutant and their roles in aging have not been determined. We hypothesized that RSKS-1-mediated translational regulation plays important roles in the significantly prolonged longevity of *daf-2 rsks-1* mutant animals. By genome-wide translational state analysis and genetic screens, we identified ribosomal protein genes and *cyc-2.1*, which encodes the worm cytochrome c ortholog, as negative regulators of longevity. Inhibition of *cyc-2.1* results in robust lifespan extension that requires both UPR^mt^ and AMPK. Intriguingly, germline is the key tissue for *cyc-2.1* to regulate longevity, and inhibition of *cyc-2.1* in the germline initiates a cell non-autonomous response that activates UPR^mt^ and AMPK in the intestine. We identified the RNA binding protein GLD-1 as a critical translational repressor of *cyc-2.1* in the germline. The synergistic lifespan extension of the *daf-2 rsks-1* mutant can be suppressed by inhibiting GLD-1 or key mediators of UPR^mt^. Therefore, the insulin-like signaling and TOR pathway-mediated tissue-specific translational repression of cytochrome c induces a cell non-autonomous mitochondrial stress response to promote longevity.

## Results

### Genome-wide translational state analysis of the significantly long-lived *daf-2 rsks-1* mutant

In order to characterize the roles of RSKS-1/S6K as a key regulator of mRNA translation in the significantly prolonged longevity of *daf-2 rsks-1*, we performed genome-wide translational state analysis via polysomal profiling coupled with RNA-Seq using wild-type N2 and *daf-2 rsks-1* mutant animals (Figure 1A). Day 2 adult animals were collected for extraction of total mRNAs and translated mRNAs (≥ 2 ribosomes / transcript) for quantification via RNA-Seq (Figure 1A). Changes in translation were determined by comparing the ratio of polysome-associated mRNAs to total mRNAs between N2 and the *daf-2 rsks-1* mutant. This is referred to as the Differential Polysome Association Ratio (DPAR). Together, we identified 167 transcripts with differential translation but no changes at total mRNA levels. Among them, 52 genes are up-regulated and 115 genes are down-regulated in the *daf-2 rsks-1* mutant (Figure 1B, Table S1). Gene Ontology (GO) enrichment analysis of biological processes revealed ‘meiotic cell cycle’, ‘cell cycle’ and ‘organelle fission’ are top three terms in the up-regulated genes, whereas ‘translation’, ‘ribosome biogenesis’ and ‘developmental process’ are the top three terms in the down-regulated genes (Figure 1C).

**Figure 1.**
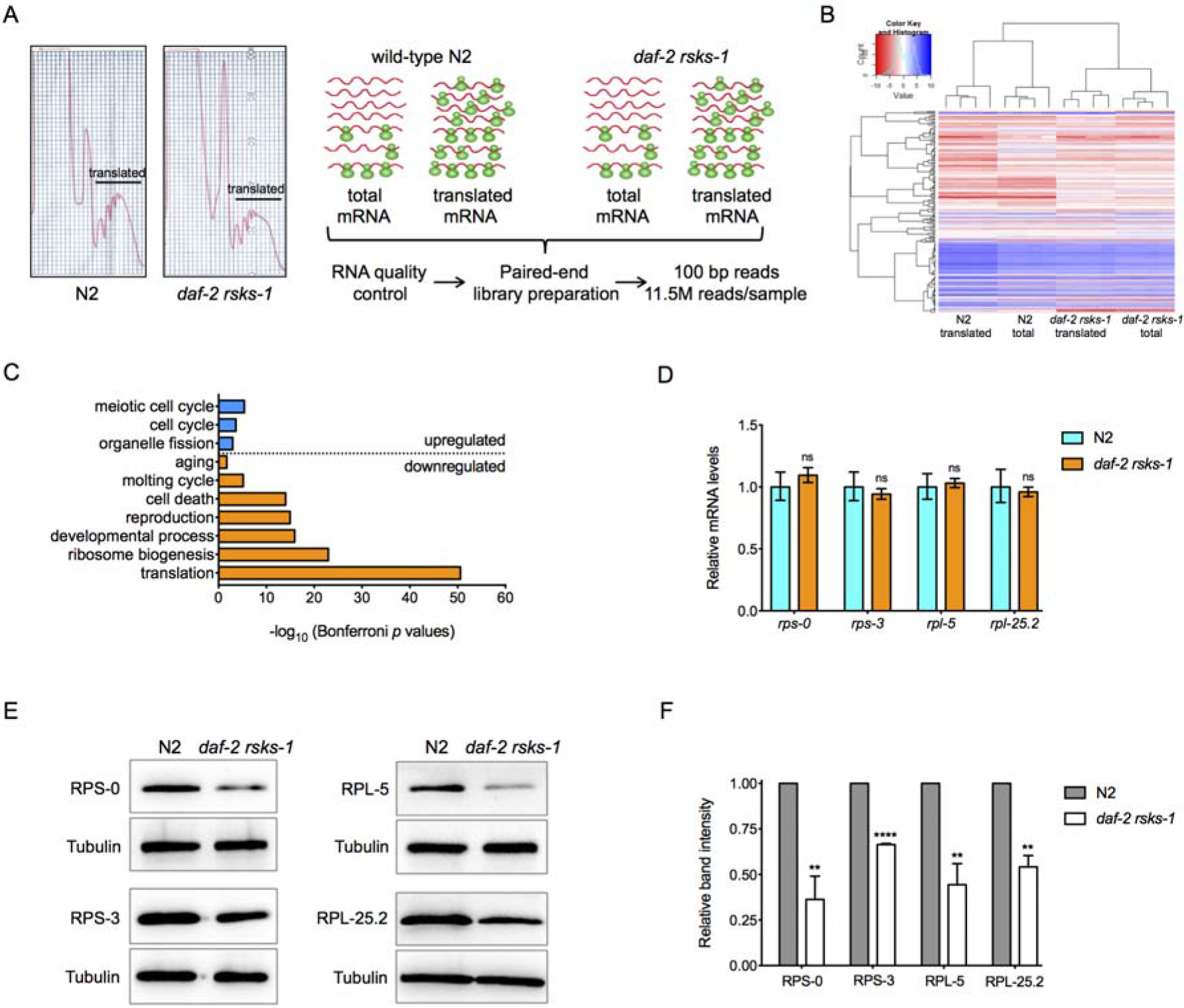
Identification of genes that are translationally regulated in the *daf-2 rsks-1* mutant. (A) Experimental design of the genome-wide translational state analysis. On the left are representative polysome profiles of wild-type N2 and the *daf-2 rsks-1* double mutant. The total and translated mRNA fractions were used for genome-wide translational state analysis as depicted by the workflow on the right. (B) Cluster of genes differentially expressed in the *daf-2 rsks-1* mutant compared to N2. (C) Top gene ontology (GO) terms for genes differentially expressed at the translational level in the *daf-2 rsks-1* mutant. (D) RT-qPCR quantification of *rps-0*, *rps-3*, *rpl-5* and *rpl-25.2* mRNA levels in N2 and the *daf-2 rsks-1* mutant (ns, *p* > 0.05, two-tailed *t* - tests). (E and F) Immunoblots (E) and quantification (F) of RPS-0, RPS-3, RPL-5, RPL-25.2 and tubulin protein levels in N2 and the *daf-2 rsks-1* mutant. Ratio of band intensity of ribosomal proteins to tubulin was normalized to N2. Data are represented as mean ± SEM based on three biological replicates (****, *p* < 0.0001, **, *p* < 0.01, two-tailed *t* - tests). See also Table S1.

To validate whether mRNAs with differential ribosome loading are indeed regulated at the mRNA translational level, we compared expression of *rps-0*, *rps-3*, *rpl-5* and *rpl-25.2* at both mRNA and protein levels between N2 and the *daf-2 rsks-1* mutant. These genes were chosen based on the availability of antibodies (Liu et al., 2018) to detect their protein products. Consistent with the RNA-Seq data (Table S1), mRNA levels of *rps-0*, *rps-3*, *rpl-5* and *rpl-25.2* show no significant changes between N2 and *daf-2 rsks-1* mutant animals (Figure 1D), whereas protein products of these genes are significantly decreased in the *daf-2 rsks-1* mutant compared to N2 (Figure 1E-F). These results indicate that polysomal profiling is a valid approach to quantitatively assess mRNA translation, which allowed us to identify differentially translated mRNAs in the *daf-2 rsks-1* mutant.

### Genes down-regulated in the *daf-2 rsks-1* mutant are enriched with lifespan determinants

Numerous studies have demonstrated that genes differentially expressed in long-lived mutants are key regulators of lifespan. We thus hypothesized that genes translationally down-regulated in the long-lived *daf-2 rsks-1* mutant are likely to be negative regulators of longevity, inhibition of which in the wild-type background could extend lifespan. Thus, we performed an RNAi-based genetic screen to individually knockdown those 115 translationally down-regulated genes in N2 to test their effects on lifespan. To facilitate the survival assays, the primary screen was performed at 25°C using the *spe-9; rrf-3* double mutant, which shows enhanced RNAi sensitivity, temperature-sensitive sterility and normal lifespan. Intriguingly, 39 out of the 115 RNAi treatments during development led to larval arrest, suggesting that genes translationally down-regulated in the *daf-2 rsks-1* mutant are enriched with developmentally essential genes. We then performed RNAi treatments against these 39 genes only during adulthood to test their lifespan phenotypes. After the re-test in the wild-type background at 20°C, we identified 24 genes, inhibition of which leads to significant lifespan extension (Table 1, Figure S1). Among them, 17 genes are essential ones that encode various ribosomal subunits. These results highlight the importance of developmentally essential genes and S6K-regulated ribosomal biogenesis in lifespan determination.

**Table 1.** List of lifespan determinant genes that are translationally repressed in the *daf-2 rsks-1* mutant

| RNAi | Lifespan (days) |  | Lifespan extension <sup>c</sup> | n <sup>d</sup> | Function | Essentiality <sup>e</sup> |
| --- | --- | --- | --- | --- | --- | --- |
|  | Mean <sup>a</sup> | Max <sup>b</sup> |  |  |  |  |
| control | 19.1 | 27 | / | 79 | / | / |
| *rpl-5* | 22.5 | 31 | 18% **** | 82 | mRNA translation | Yes |
| *rpl-6* | 23.2 | 35 | 22% **** | 76 | mRNA translation | Yes |
| *rpl-7* | 23.5 | 33 | 23% **** | 85 | mRNA translation | Yes |
| *rpl-13* | 23.1 | 33 | 21% **** | 89 | mRNA translation | Yes |
| *rpl-14* | 22.1 | 31 | 16% **** | 86 | mRNA translation | Yes |
| *rpl-15* | 21.6 | 35 | 13% **** | 73 | mRNA translation | Yes |
| *rpl-18* | 22.1 | 33 | 16% **** | 74 | mRNA translation | Yes |
| *rpl-25.2* | 22.1 | 29 | 16% **** | 87 | mRNA translation | Yes |
| *rpl-30* | 22.3 | 35 | 17% **** | 74 | mRNA translation | Yes |
| *rpl-33* | 23.6 | 37 | 24% **** | 82 | mRNA translation | Yes |
| *rpl-35* | 22.1 | 31 | 16% **** | 78 | mRNA translation | Yes |
| *rpl-36* | 23.8 | 33 | 25% **** | 73 | mRNA translation | Yes |
| *rps-3* | 22.8 | 33 | 20% **** | 77 | mRNA translation | Yes |
| *rps-12* | 22.3 | 33 | 17% **** | 63 | mRNA translation | Yes |
| *rps-20* | 22.2 | 35 | 17% **** | 80 | mRNA translation | Yes |
| *rps-26* | 21.2 | 31 | 11% *** | 66 | mRNA translation | Yes |
| *rps-30* | 22.7 | 33 | 17% **** | 77 | mRNA translation | Yes |
| *cytb-5.2* | 22.1 | 33 | 16% **** | 73 | metabolism | No |
| *cyc-2.1* | 33.8 | 49 | 77% **** | 78 | mitochondria | No |
| *sym-1* | 20.9 | 31 | 9% ** | 57 | development | No |
| *ncam-1* | 21.4 | 31 | 12% *** | 70 | development | No |
| *pqn-48* | 20.6 | 29 | 8% * | 69 | protein modification | No |
| *pqn-65* | 20.8 | 29 | 9% ** | 70 | unknown | No |
| *T10B5.7* | 22.0 | 31 | 15% **** | 75 | unknown | No |

<sup>a</sup> Mean lifespan.
<sup>b</sup> Maximum lifespan.
<sup>c</sup> Mean lifespan extension compared to the control RNAi treated animals. \*\*\*\*, $p < 0.0001$ ; \*\*\*, $p < 0.001$ ; \*\*, $p < 0.01$ ; \*, $p < 0.05$ (log-rank tests).
<sup>d</sup> Numbers of animals counted.
<sup>e</sup> Essential genes are those, RNAi knocking-down of which during development led to larval arrest. Lifespan assays of these genes were performed with RNAi treatments only during adulthood.
See also Figure S1.

### Inhibition of CYC-2.1 results in robust lifespan extension that requires UPR^mt^ and AMPK

*cyc-2.1*, which encodes a highly conserved cytochrome c ortholog, showed the strongest lifespan extension upon RNAi inhibition (Table 1, Figure S1). Knockdown of *cyc-2.1* significantly extends lifespan not only in wild-type animals but also in the long-lived *daf-2* single mutant (Figure 2A, Table S2).

**Figure 2.**
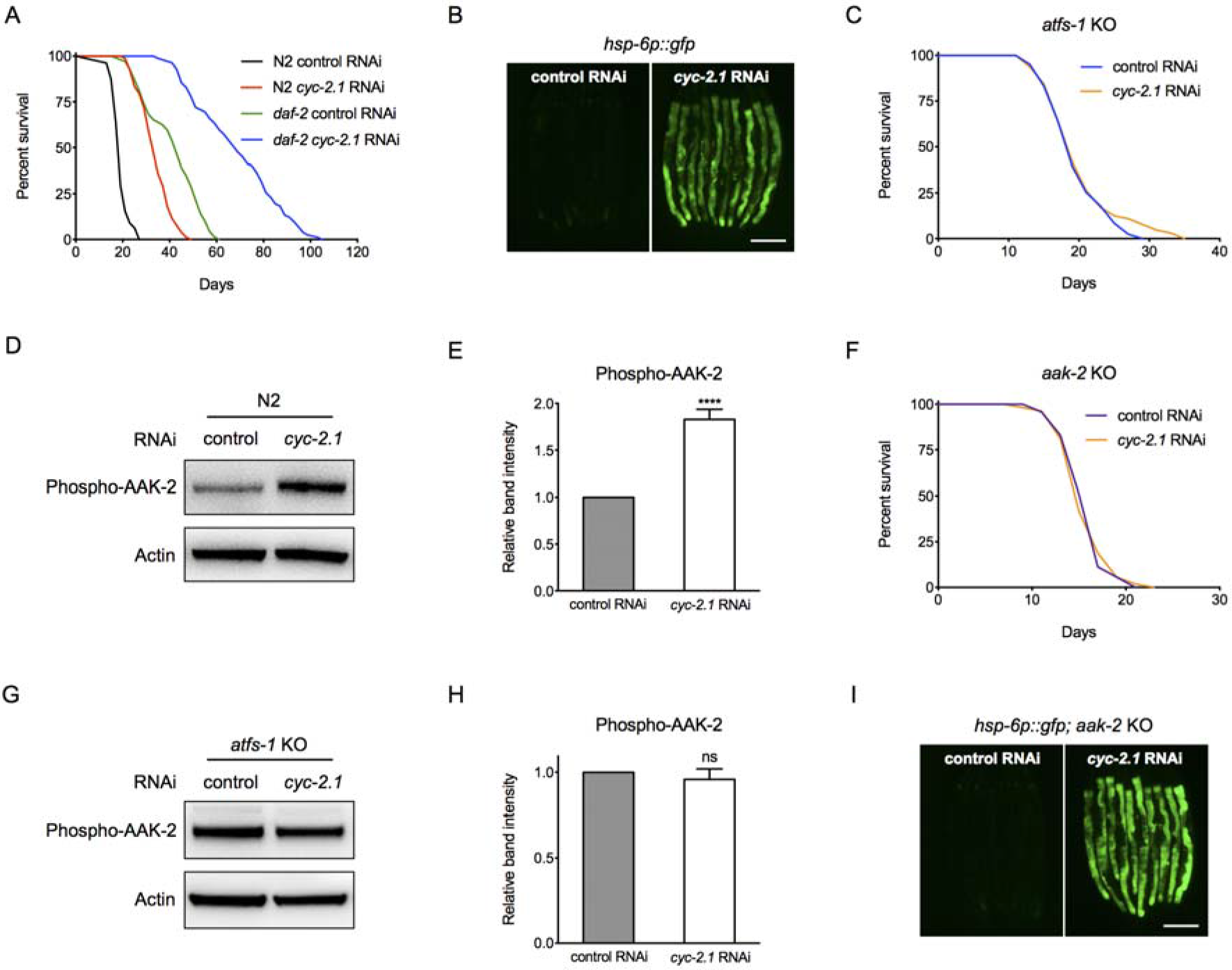
Knockdown of *cyc-2.1* significantly extends lifespan by activating UPR^mt^ and AMPK. (A) Survival curves of N2 and the *daf-2* mutant treated with either control or *cyc-2.1* RNAi (*p* < 0.0001, log-rank tests). (B) Representative photographs of *hsp-6p::gfp* expression in animals treated with either control or *cyc-2.1* RNAi. (C) Survival curves of the *atfs-1* KO mutant treated with control or *cyc-2.1* RNAi (*p* = 0.2985, log-rank test). (D and E) Immunoblots (D) and quantification (E) of phospho-AAK-2 (AMPKα) and actin in N2 animals treated with either control or *cyc-2.1* RNAi. Ratio of band intensity of phospo-AAK-2 to actin was normalized to the control RNAi treated animals. Data are represented as mean ± SEM based on six biological replicates (****, *p* < 0.0001, two-tailed *t* - test). (F) Survival curves of the *aak-2* KO mutant treated with control or *cyc-2.1* RNAi (*p* = 0.9647, log-rank test). (G and H) Immunoblots (G) and quantification (H) of phospho-AAK-2 and actin in the *atfs-1* KO mutant treated with either control or *cyc-2.1* RNAi. Data are represented as mean ± SEM based on four biological replicates (ns, *p* = 0.5239, two-tailed *t* - test). (I) Representative photographs of *hsp-6p::gfp* expression in *aak-2* KO mutant animals treated with either control or *cyc-2.1* RNAi. Scale bars, 200 μm. See also Figure S2 and Table S2.

Cytochrome c functions in the mitochondrial electron transport chain (ETC) by transferring electrons from complex III to complex IV. Previous studies demonstrated that inhibition of certain mitochondrial ETC components extends lifespan, and the underlying mechanisms involve CEP-1/p53 (Baruah et al., 2014), the intrinsic apoptosis pathway (Yee et al., 2014) and UPR^mt^ (Dillin et al., 2002; Durieux et al., 2011; Lee et al., 2002). Mutations in the *C. elegans* p53 ortholog CEP-1 or a key component of the apoptosis pathway CED-4 significantly suppress the lifespan extension by mitochondrial ETC mutants (Baruah et al., 2014; Yee et al., 2014). However, *cyc-2.1* RNAi treatment significantly extends lifespan in null mutants of *cep-1* or *ced-4* (Figure S2), suggesting a potential novel mechanism. To examine the effect of *cyc-2.1* knockdown on the mitochondrial stress response, we applied *cyc-2.1* RNAi to transgenic animals carrying a *gfp* reporter driven by the *hsp-6* promoter, which has been widely used to monitor UPR^mt^ activation. Inhibition of *cyc-2.1* significantly activates *hsp-6p::gfp* expression (Figure 2B). Previous studies have identified ATFS-1 as one of the major transcription factors that mediate the UPR^mt^ activation (Nargund et al., 2012). *cyc-2.1* RNAi-induced longevity phenotype is completely suppressed by a deletion mutant of *atfs-1* (Figure 2C, Table S2). Therefore, activated mitochondrial stress response plays an essential role in cytochrome c knockdown-induced lifespan extension.

It has been reported that paraquat, a ROS (reactive oxygen species) generator, and hypomorphic mutations in mitochondrial ETC genes such as *isp-1*, extend *C. elegans* lifespan by activating AMPK (Hwang et al., 2014). The synergistic lifespan extension by *daf-2 rsks-1* also requires AMPK (Chen et al., 2013). We then performed immunoblots to measure the levels of phosphorylated AAK-2 (AMPKα), which serve as an indicator of AMPK activation, in control or *cyc-2.1* RNAi treated animals. Knockdown of *cyc-2.1* significantly increases phospho-AAK-2 levels compared to the control (Figure 2 D-E). Consistently, *cyc-2.1* RNAi fails to extend lifespan of the *aak-2* deletion mutant (Figure 2F, Table S2). Therefore, inhibition of *cyc-2.1* activates UPR^mt^ and AMPK to extend lifespan.

To characterize the relationship between UPR^mt^ and AMPK in *cyc-2.1* knockdown-induced lifespan extension, we first measured phospho-AAK-2 levels in the *atfs-1* null mutant treated with either control or *cyc-2.1* RNAi. Immunoblots and quantification results indicate that unlike in the N2 background, *cyc-2.1* RNAi does not increase phospho-AAK-2 levels in the *atfs-1* mutant (Figure 2 G-H). We then crossed the *hsp-6* promoter::*gfp* reporter into the *aak-2* deletion mutant. In the absence of AAK-2, *cyc-2.1* RNAi still significantly activates the *hsp-6::gfp* reporter (Figure 2I). Together, these results demonstrate that AMPK functions downstream of UPR^mt^ activation to promote lifespan extension produced by the *cyc-2.1* RNAi treatment.

### Tissue-specific knockdown of *cyc-2.1* non-autonomously activates UPR^mt^ and AMPK to extend lifespan

Previous studies demonstrated that the activation of UPR^mt^ can be achieved by tissue-specific perturbation of mitochondrial ETC in a cell non-autonomous manner (Durieux et al., 2011). To determine the key tissues, in which *cyc-2.1* functions to regulate lifespan as well as the activation of UPR^mt^ and AMPK, we performed tissue-specific RNAi experiments to knockdown *cyc-2.1* in the germline, intestine, epidermis and muscles, respectively. Spatially restricted RNAi knockdown was achieved by tissue-specific promoters-driving transgene rescue of mutations in *rde-1*, which is essential for the RNAi machinery to be functional (Espelt et al., 2005; Qadota et al., 2007; Zou et al., 2018). Knockdown of *cyc-2.1* in the germline significantly extends lifespan by 27% (Figure 3A, Figure S3A, Table S2). However, knockdown of *cyc-2.1* in the intestine has no effect on lifespan (Figure 3A, Figure S3B, Table S2). Inhibition of *cyc-2.1* by RNAi in the epidermis or muscles does not significantly affect lifespan (Figure 3A, Figure S3 C-D, Table S2). The importance of germline in *cyc-2.1* RNAi-mediated lifespan extension is further supported by the evidence that knockdown of *cyc-2.1* in the germline-less *glp-4(ts)* mutant (Beanan and Strome, 1992) only extends lifespan by 9% (Figure S2E). Together, these results demonstrate that germline is the key tissue in which *cyc-2.1* knockdown significantly extends lifespan.

**Figure 3.**
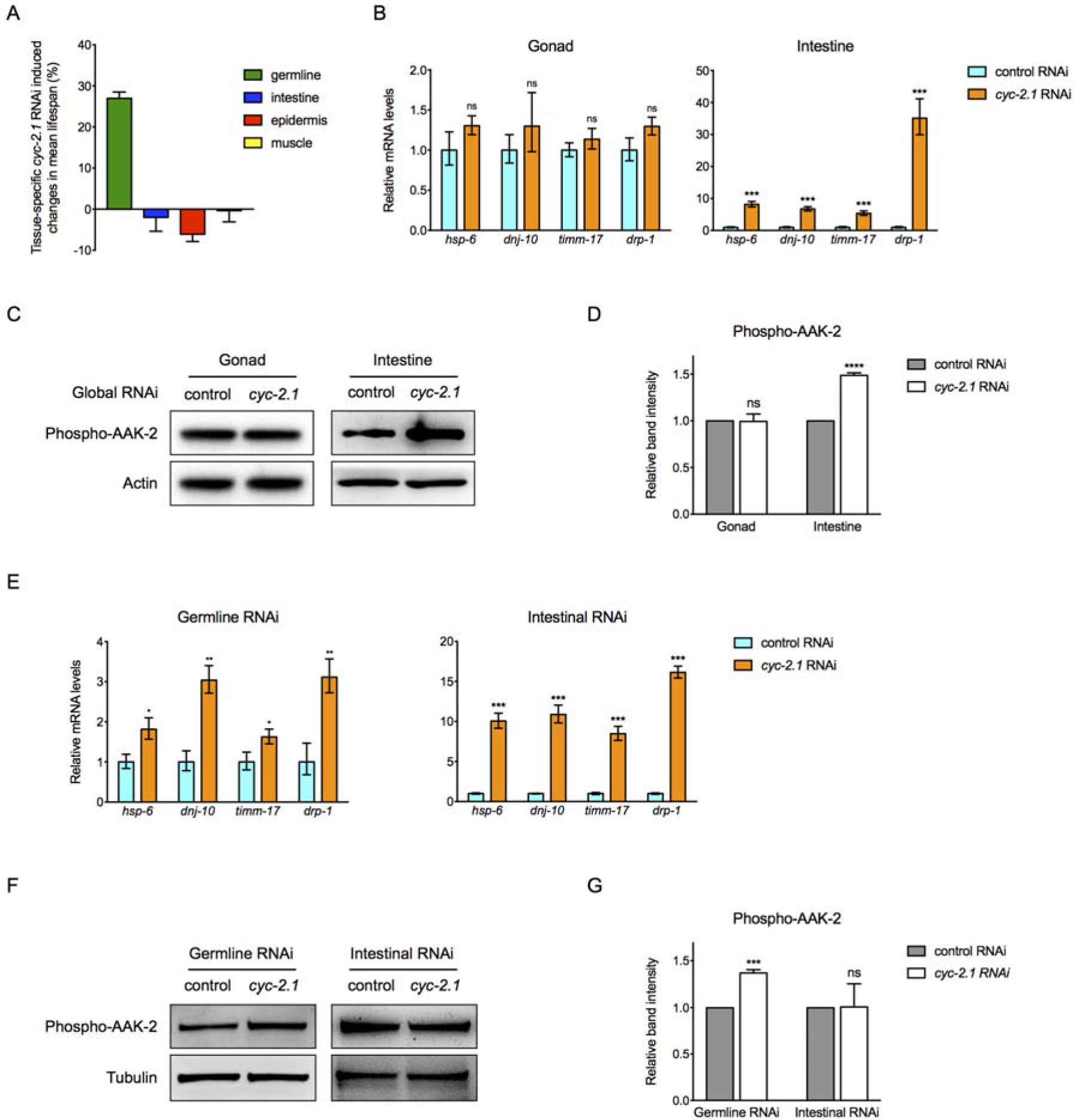
Inhibition of *cyc-2.1* in the germline extends lifespan by cell-non-autonomous activation of UPR^mt^ and AMPK in the intestine. (A) Tissue-specific *cyc-2.1* RNAi-induced changes in the mean lifespan relative to control RNAi-treated animals. Data are represented as mean ± SEM based on three biological replicates. (B) RT-qPCR quantification of UPR^mt^ markers *hsp-6*, *dnj-10*, *timm-17* and *drp-1* using RNAs extracted from dissected gonadal and intestinal tissues, respectively (***, *p* < 0.001; ns, *p* > 0.05, two-tailed *t* - tests). (C and D) Immunoblots (C) and quantification (D) of phospho-AAK-2 and actin using proteins extracted from dissected gonadal and intestinal tissues, respectively. Data are represented as mean ± SEM based on three biological replicates (ns, *p* = 0.9475, ****, *p* < 0.0001, two-tailed *t* - tests). (E) RT-qPCR quantification of UPR^mt^ markers *hsp-6*, *dnj-10*, *timm-17* and *drp-1* using RNAs extracted from dissected intestinal tissues with germline- and intestine-specific *cyc-2.1* RNAi treatment, respectively (***, *p* < 0.001; **, *p* < 0.01, two-tailed *t* - tests). (F and G) Immunoblots (F) and quantification (G) of phospho-AAK-2 and actin using proteins extracted from dissected intestinal tissues treated with germline- or intestine-specific control vs. *cyc-2.1* RNAi. Data are represented as mean ± SEM based on three biological replicates (****, *p* < 0.0001, ns, *p* = 0.5504, two-tailed *t* - tests). See also Figure S3–5 and Table S2.

To better understand whether *cyc-2.1* RNAi activates UPR^mt^ in a tissue-specific manner, we performed qPCR experiments to measure mRNA levels of several ATFS-1 direct target genes including *hsp-6*, *dnj-10*, *timm-17* and *drp-1*, transcription levels of which are elevated upon mitochondrial ETC perturbation (Nargund et al., 2012; 2015). Gonadal and intestinal tissues were dissected from wild-type animals treated with either control or *cyc-2.1* RNAi for RT-qPCR assays. Since the gonad is 95% germline and 5% somatic gonad, and the germline and somatic gonad cannot be further dissected, we used the gonadal tissue as the proxy for the germline in subsequent experiments. Surprisingly, *cyc-2.1* RNAi treatment significantly activates UPR^mt^ markers in the intestine, but not in the germline (Figure 3B), although the latter is the tissue in which *cyc-2.1* functions to regulate lifespan. Consistently, global *cyc-2.1* RNAi treatment significantly activates AMPK in the intestine but not in the gonad (Figure 3 C-D). Although both germline and intestinal-specific *cyc-2.1* RNAi treatments significantly activate UPR^mt^ markers in the intestine (Figure 3E), only the germline but not intestinal *cyc-2.1* RNAi treatment led to increased phospho-AAK-2 levels in the intestine (Figure 3 F-G). Therefore, knockdown of *cyc-2.1* in the germline causes non-autonomous activation of UPR^mt^ and AMPK in the intestine, suggesting the existence of a germline to intestine signaling to regulate UPR^mt^, AMPK and lifespan.

### Tissue-specific activation of UPR^mt^ contributes to the significantly prolonged longevity of the *daf-2 rsks-1* mutant

*cyc-2.1* was identified as one of the genes that show significantly decreased ribosomal loading in the *daf-2 rsks-1* mutant compared to the wild-type N2 from the translational profiling analysis (Table S1). To examine whether CYC-2.1 is repressed at the protein level in the *daf-2 rsks-1* mutant, we used CRISPR/Cas9-based genome editing approach to knock in a 3×FLAG tag to the C terminus of CYC-2.1 for immunoblots using anti-FLAG antibodies. CYC-2.1 protein levels are significantly reduced in the gonad but not intestine of the *daf-2 rsks-1* mutant (Figure 4 A-B). RT-qPCR measurement of UPR^mt^ markers using dissected tissues demonstrates that the *daf-2 rsks-1* mutant has UPR^mt^ activation in the intestine, but not in the gonad (Figure 4C).

**Figure 4.**
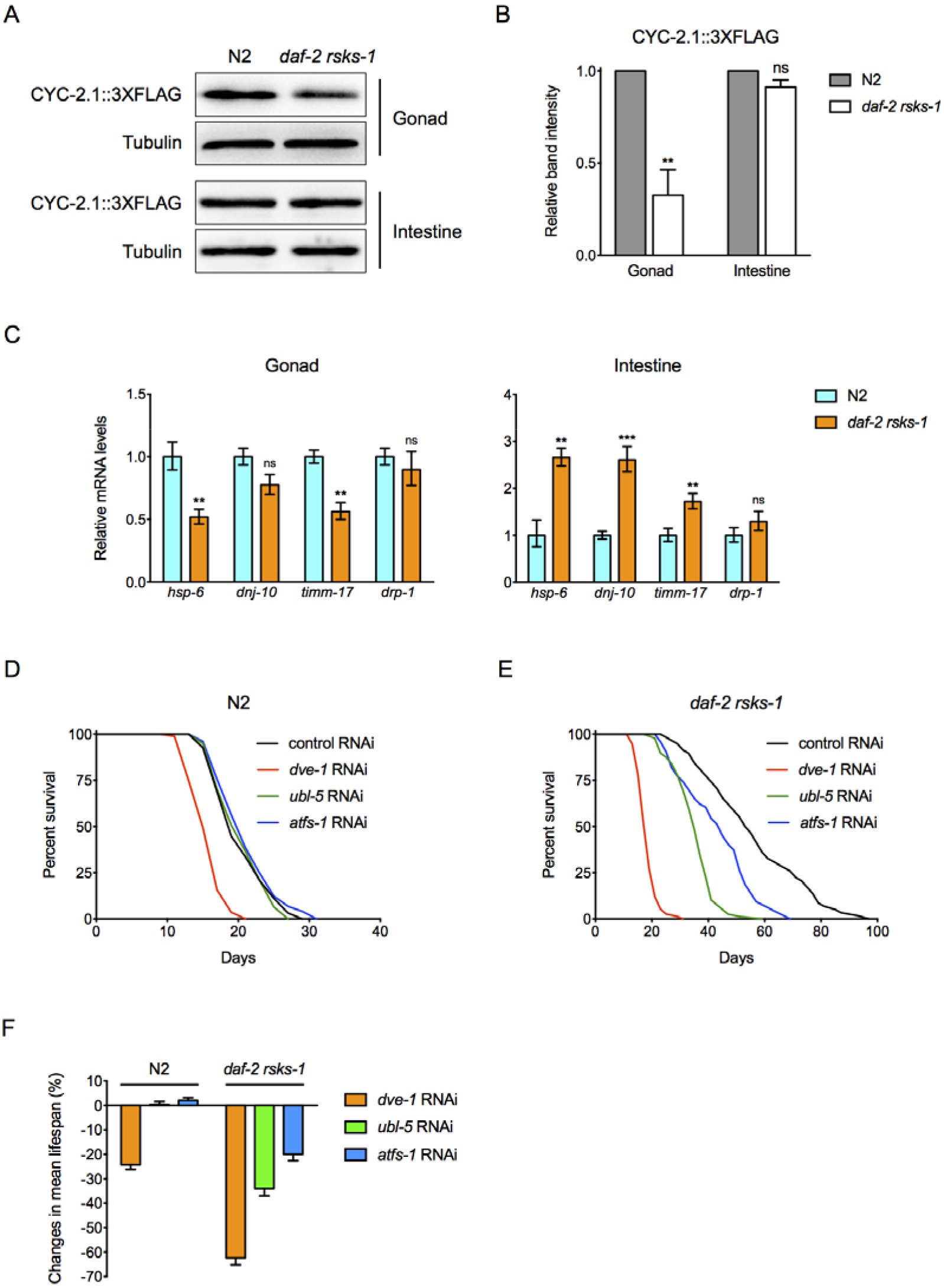
Translational repression of CYC-2.1 in the germline and non-autonomous activation of UPR^mt^ in the intestine play an important role in regulating the synergistic lifespan extension by *daf-2 rsks-1*. (A and B) Immunoblots (A) and quantification (B) of CYC-2.1::3×FLAG and tubulin using proteins extracted from dissected gonadal and intestinal tissues, respectively. Data are represented as mean ± SEM based on three biological replicates (**, *p* < 0.01, ns, *p* = 0.0880, two-tailed *t* - tests). (C) RT-qPCR quantification of UPR^mt^ markers *hsp-6*, *dnj-10*, *timm-17* and *drp-1* using RNAs extracted from dissected gonadal and intestinal tissues, respectively (**, *p* < 0.01; ***, *p* < 0.001; ns, *p* > 0.05, two-tailed *t* - tests). (D and E) Survival curves of N2 (D) and the *daf-2 rsks-1* mutant (E) treated with *dve-1*, *ubl-5* or *atfs-1* RNAi, respectively. (E) Changes in mean lifespan relative to control RNAi-treated animals by knockdown of *dve-1*, *ubl-5* or *atfs-1* in N2 or *daf-2 rsks-1* mutant. Data are represented as mean ± SEM based on three biological replicates. See also Table S2.

Previous studies showed that transcriptional factors DVE-1 and ATFS-1, as well as UBL-5, a small ubiquitin-like factor that serves as the co-factor for DVE-1, are the key mediators of UPR^mt^ (Haynes et al., 2007; Nargund et al., 2012). Inhibition of *dve-1* by RNAi shortens N2 lifespan, whereas *ubl-5* and *atfs-1* RNAi treatments do not affect N2 lifespan (Fig. 4D, F, Table S2). The prolonged longevity of *daf-2 rsks-1* mutant animals can be significantly decreased by *dve-1*, *ubl-5* or *atfs-1* RNAi (Figure 4E, F, Table S2). Together, the *daf-2 rsks-1* mutant shows germline reduction of CYC-2.1 and intestinal activation of UPR^mt^, which is required for the significantly prolonged longevity of the *daf-2 rsks-1* mutant.

### GLD-1-mediated translational regulation of CYC-2.1 is important for the UPR^mt^ activation and significantly prolonged longevity in the *daf-2 rsks-1* mutant

Specific translational regulations in many cases are mediated by RNA binding proteins and their association with 5′- or 3′-UTRs (untranslated regions). GLD-1, a K homology (KH) RNA binding protein, is regulated by the GLP-1/Notch signaling to negatively regulate target genes’ translation in the germline (Kimble and Crittenden, 2005). Our previous studies showed that a *glp-1* gain-of-function mutation, which decreases GLD-1 expression, suppresses the significantly extended lifespan of *daf-2 rsks-1* (Chen et al., 2013). Our genome-wide translational state analysis indicates that *gld-1* mRNAs have elevated ribosomal loading in the *daf-2 rsks-1* mutant (Table S1). Together, these results suggest that the germline-specific translational repressor GLD-1 might be involved in regulating CYC-2.1 protein levels in the germline of the *daf-2 rsks-1* mutant.

To test whether GLD-1 is up-regulated in the *daf-2 rsks-1* mutant, we performed CRISPR/Cas9 based genome editing experiments to knock-in the mKate2 red fluorescent protein coding sequence to the 3′ of *gld-1*. Compared to the wild-type control, GLD-1::mKate2 protein levels are significantly increased in the germline of *daf-2 rsks-1* mutant animals (Figure 5 A-B). To examine whether GLD-1 is involved in the translational regulation of *cyc-2.1*, we applied *gld-1* RNAi to the *daf-2 rsks-1* mutant and found that CYC-2.1 protein levels are significantly elevated compared to the control RNAi treated animals (Figure 5 C-D). Consistently, knockdown of *gld-1* also significantly decreases UPR^mt^ activation in the intestine of *daf-2 rsks-1* mutant animals (Figure 5E). Furthermore, inhibition of *gld-1* by RNAi significantly decreases lifespan of the *daf-2 rsks-1* mutant (Figure 5F). Together, these results demonstrate that GLD-1 functions as a translational repressor of *cyc-2.1* in the germline to non-autonomously activate UPR^mt^ and extend lifespan in the *daf-2 rsks-1* mutant.

**Figure 5.**
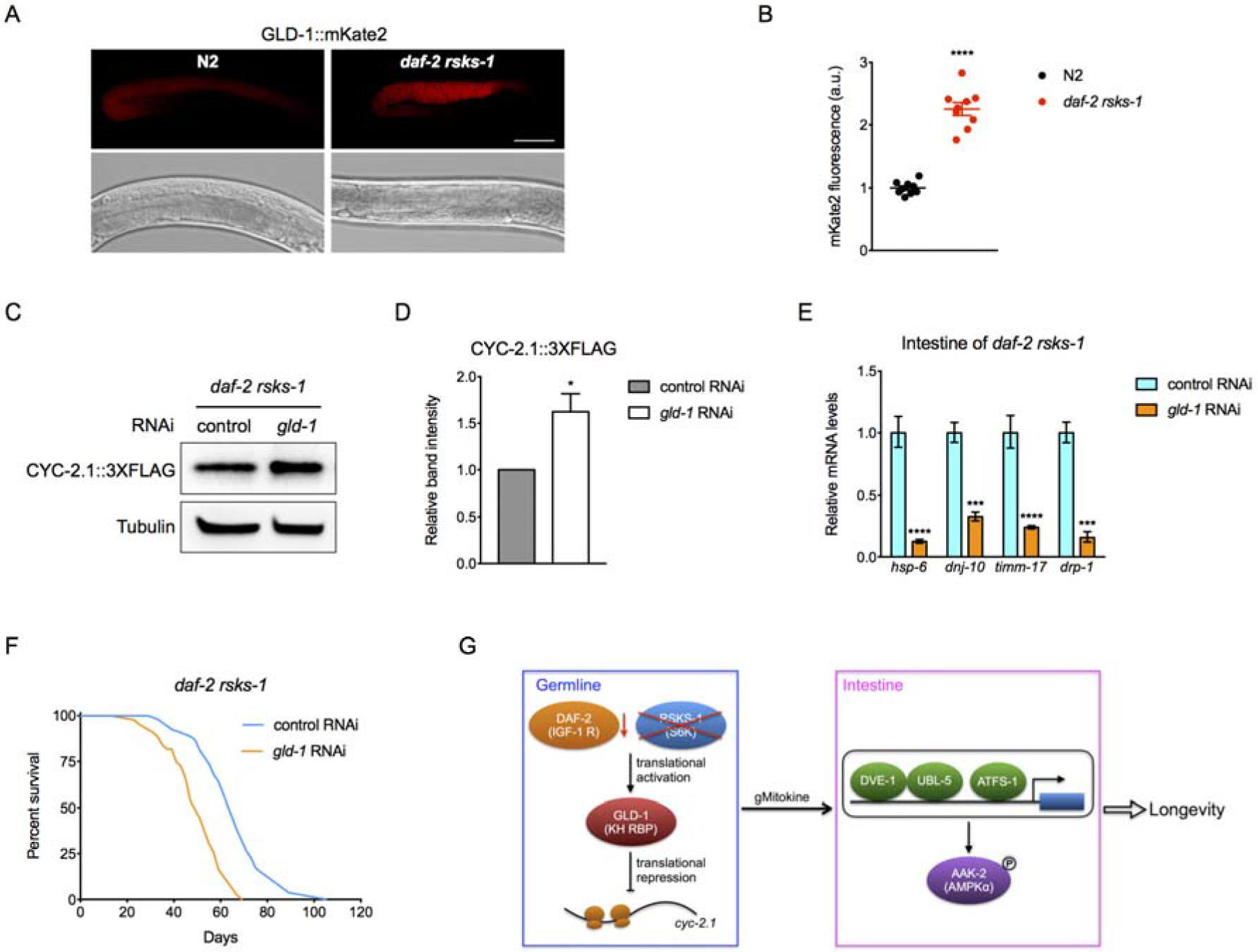
Translational repression of *cyc-2.1* by GLD-1 contributes to the UPR^mt^ activation and lifespan extension in the *daf-2 rsks-1* mutant. (A and B) Representative photographs (A) and quantification (B) of GLD-1::mKate2 expression in the germline of N2 and *daf-2 rsks-1* mutant animals. Data are represented as mean ± SEM (****, *p* < 0.0001, two-tailed *t* - tests). Scale bar, 50 μm. (C and D) Immunoblots (C) and quantification (D) of CYC-2.1::3×FLAG and tubulin in *daf-2 rsks-1* mutant animals treated with either control or *gld-1* RNAi. Data are represented as mean ± SEM based on three biological replicates (*, *p* < 0.05, two-tailed *t* - test). (E) RT-qPCR quantification of UPR^mt^ markers *hsp-6*, *dnj-10*, *timm-17* and *drp-1* using RNAs extracted from dissected intestinal tissues of the *daf-2 rsks-1* mutant treated with the control or *gld-1* RNAi (***, *p* < 0.001; ****, *p* < 0.0001, two-tailed *t* - tests). (F) Survival curves of *daf-2 rsks-1* mutant animals treated with either control or *gld-1* RNAi (*p* < 0.0001, log-rank test). (G) A model depicting the translational repression of CYC-2.1 by GLD-1 in the germline that non-autonomously activates UPR^mt^ and AMPK in the intestine via the germline produced mitokine (gMitokine), which leads to significantly extended lifespan in the *daf-2 rsks-1* mutant. See also Table S2.

In summary, we have performed genome-wide translational state analysis via polysomal profiling coupled with RNA-Seq and identified genes that are regulated at the translational level in the significantly long-lived *daf-2 rsks-1* mutant. Mechanistically, we demonstrated that GLD-1, an RNA binding protein, is up-regulated at the protein level in the germline of *daf-2 rsks-1* mutant. Elevated GLD-1 leads to translational repression of CYC-2.1, the worm cytochrome c ortholog, in the germline, which activates UPR^mt^ and AMPK in the intestine and significantly extends lifespan (Figure 5G). These results highlight the importance of translational regulation of a highly conserved mitochondrial gene in the significantly prolonged longevity exerted by inhibiting both the insulin-like signaling and S6K.

## Discussion

The highly conserved insulin/insulin-like signaling (IIS) and TOR pathway play an important role in aging across species (Fontana et al., 2010; Kenyon, 2010). To examine how these pathways interact with each other to modulate aging, we previously constructed a *daf-2 rsks-1* double mutant that carries loss-of-function mutations in the DAF-2/IGF-1 receptor and TOR effector RSKS-1/S6K. The double mutant shows a synergistic rather than additive effect on longevity, suggesting active interactions between these two important aging-related pathways. Functional genomics studies via transcriptome profiling helped to identify AMPK-mediated positive feedback regulation of the DAF-16/FOXO transcription factor mechanism in the *daf-2 rsks-1* mutant (Chen et al., 2013). However, RSKS-1, which serves as a key regulator of mRNA translation, has not been well characterized for its roles in the significantly extended longevity of *daf-2 rsks-1* mutant animals.

It has been well documented that inhibition of translation delays aging (Hansen et al., 2007; Kapahi et al., 2010; 2004; Pan et al., 2007; Rogers et al., 2011). One hypothesis is that reduced global translation helps organisms to maintain better protein homeostasis, dysregulation of which leads to aging and age-related pathologies. A not mutually exclusive hypothesis suggests that the anti-aging effect is achieved by the translational regulation of key modulators of aging. We reasoned that identification of genes that are translationally regulated in the *daf-2 rsks-1* mutant and characterization of the translational regulation should help to gain better mechanistic insights into the significantly extended longevity of the *daf-2 rsks-1* double mutant.

Through genome-wide translational state analysis and genetic screens, we identified negative regulators of longevity from the 115 genes that are translationally down-regulated in the *daf-2 rsks-1* mutant. One intriguing observation was the lifespan determinants are enriched with developmentally essential genes (71%). Inhibition of these genes during development causes larval arrest, whereas RNAi knockdown only during adulthood leads to significant lifespan extension (Table 1, Figure S1). These results support the antagonistic pleiotropy theory of aging, which proposes that aging is adaptive since natural selection favors genes that confer benefits during development but cause deleterious effects later in life (Williams, 1957). Thus, inhibition of developmentally essential genes during adulthood might extend lifespan (Chen et al., 2007; Curran and Ruvkun, 2007). All of the developmentally essential, lifespan determinant genes encode various ribosomal subunits (Table 1), which is consistent with previous studies showing that TOR-mediated ribosomal biogenesis plays an important role in aging (Steffen et al., 2008).

Among the lifespan regulators that we identified, knockdown of *cyc-2.1*, which encodes the *C. elegans* cytochrome c ortholog that transfers electrons from mitochondrial ETC complex III to complex IV, results in the most significant lifespan extension (Table 1, Figure 2A, Figure S1F). Inhibition of genes encoding components of mitochondrial respiratory complexes extends lifespan (Dillin et al., 2002; Houtkooper et al., 2014; Lee et al., 2002), and the underlying mechanisms involve UPR^mt^ (Durieux et al., 2011), CEP-1/p53 (Baruah et al., 2014) and the intrinsic apoptosis pathway (Yee et al., 2014). CEP-1/p53 and CED-4 from the intrinsic apoptosis pathway have been identified as key mediators of mitochondrial ETC gene perturbation-induced longevity (Baruah et al., 2014; Yee et al., 2014). However, *cyc-2.1* RNAi can still significantly extend lifespan in null mutants of *cep-1*, *ced-13* and *ced-4* (Figure S3). Inhibition of *cco-1*, which encodes the cytochrome c oxidase-1 subunit Vb/COX4, significantly extends lifespan (Dillin et al., 2002). ATFS-1 is partially required for *cco-1* RNAi-induced lifespan extension (Figure S4A), whereas the prolonged longevity by *cyc-2.1* RNAi can be completely suppressed by an *atfs-1* KO mutant (Figure 2C, Table S2). Unlike *cyc-2.1*, intestine-specific knockdown of *cco-1* is sufficient to significantly extend lifespan (Figure S4B). Taken together, these results suggest that *cyc-2.1* might function through novel mechanisms to regulate lifespan.

Previous studies showed that perturbation of mitochondrial ETC in *C. elegans* neurons provokes a pro-longevity signal to non-autonomously activate UPR^mt^ in the intestine (Berendzen et al., 2016; Durieux et al., 2011; Shao et al., 2016). In this study, we found that inhibition of *cyc-2.1* in the germline non-autonomously activates UPR^mt^ and AMPK in the intestine. Although intestine-specific *cyc-2.1* RNAi also activates UPR^mt^, it fails to activate AMPK and extend lifespan (Figure 3A, E-G, Figure S3B, Table S2). These results indicate that activation of UPR^mt^ is required but not sufficient for the lifespan extension produced by *cyc-2.1* knockdown. Since there is no direct contact between the germline and intestine in *C. elegans*, we propose that inhibition of cytochrome c in the germline might produce an endocrine-like signaling, named gMitokine (germline-produced mitokine), to activate UPR^mt^ in the distal tissue. Previous studies on communications between the *C. elegans* gonad and intestine largely focused on the transportation of intestinally produced yolk proteins and associated lipids to oocytes. This process is mediated by secretion and endocytosis (Grant, 2006). It will be important to understand whether the material transportation from the germline to intestine is also mediated by similar mechanisms. Further characterization of this germline to intestine signal transduction process, especially identifying the germline-produced mitokine and its downstream effectors, will help to better understand UPR^mt^-mediated anti-aging mechanisms.

The germline tissue plays a very important role in *C. elegans* aging. Germline-less worms showed significant lifespan extension that is dependent on the DAF-16/FOXO transcription factor and DAF-12/nuclear hormone receptor. The prolonged longevity of germline-less animals is not due to sterility, rather caused by diminished pro-aging signaling from the germline (Berman and Kenyon, 2006; Hsin and Kenyon, 1999). However, the molecular identity of aging signals produced by germ cells has not been determined. We speculate that *cyc-2.1* knockdown and germline deficiency function through different mechanisms to regulate lifespan because unlike the long-lived germline-less animals, the prolonged longevity of animals treated with *cyc-2.1* RNAi is independent of DAF-16 or DAF-12 (Figure S5).

It has also been shown that the germline is the key tissue for RSKS-1 to regulate lifespan. Knockdown of *rsks-1* in the germline of *daf-2* mutant produces a synergistic effect on longevity (Chen et al., 2013). H3K3me3 deficiency-induced lifespan extension is also mediated by down-regulation of RSKS-1 in the germline, which leads to increased accumulation of mono-unsaturated fatty acids (MUFAs) in the distal intestine tissue (Han et al., 2017). It will be interesting to test whether inhibition of *cyc-2.1* in the germline will affect lipid metabolism in the intestine, and if so, whether UPR^mt^ is involved in this process in future studies.

In conclusion, genome-wide translational state analysis allowed us to identify a series of translational regulation of lifespan determinant genes in the significantly long-lived *daf-2 rsks-1* mutant. Functional studies revealed that an RNA-binding protein GLD-1 mediated translational suppression of cytochrome c in the germline leads to non-autonomous activation of UPR^mt^ and AMPK in the intestine, which is indispensable for the significantly extended lifespan. In the future, it will be important to understand how the translational regulation is achieved, and how the germline signals the distal metabolic tissue upon mitochondrial perturbation at the molecular level.

## Acknowledgments

We thank members of the Chen lab for discussions and comments on the manuscript, and Drs. Shiqing Cai, Mengqiu Dong, Arjumand Ghazi, Cole Haynes, Ying Liu, Yubing Liu, Shohei Mitani, Ye Tian, Xiaochen Wang and Zhiping Wang for worm strains, plasmids, antibodies and unpublished results. Some strains were provided by the CGC, which is funded by NIH Office of Research Infrastructure Programs (P40 OD010440), and some strains were provided by the Japanese National BioResource Project. D.C. is supported by the National Natural Science Foundation of China (31471379, 31671527) and Natural Science Foundation of Jiangsu, China (BK20141316).

## Author contributions

J.L., D.W., and X.Z. performed experiments. J.R. performed experiments and bioinformatics analysis. L.Z., Z.W., C.Y. and Z.W. provided technical assistance. D.C., A.N.R and P.K. conceived the project, designed experiments, and wrote the paper.

## CONTACT FOR REAGENT AND RESOURCE SHARING

Further information and requests for reagents may be directed to and will be fulfilled by Di Chen.

## EXPERIMENTAL MODEL AND SUBJECT DETAILS

### *C. elegans* Strains and Maintenance

The following *C. elegans* strains used in this study were obtained from the Caenorhabditis Genome Center: Bristol (N2) strain as the wild-type strain, CB1370 *daf-2(e1370) III*, CF1038 *daf-16(mu86) I*, TJ1060 *spe-9(hc88) I; rrf-3(b26) II*, VP303 *rde-1(ne219) V; kbIs7[nhx-2p::rde-1(+) + rol-6]*, WM118 *rde-1(ne300) V; neIs9[myo-3p::HA::rde-1(+) + rol-6] X*, NR222 *rde-1(ne219) V; kzIs9[lin-26p::NLS::GFP + lin-26p::rde-1(+) + rol-6]*, SJ4100 *zcIs13[Phsp-6::gfp] V*, MT2547 *ced-4(n1162) III*, TJ1 *cep-1(gk138) I*, AA86 *daf-12(rh61rh411) X*. The following strain used in this study was obtained from the National BioResource Project: *atfs-1(tm4525) V*. The following strains used in this study were generated in D.C. and P.K. labs: DCL124 *zcIs13[Phsp-6::gfp] V; aak-2(ok524) X*, DCL178 *cyc-2.1[mkc6(cyc-2.1::3×flag)] IV*, DCL198 *daf-2(e1370) rsks-1(ok1255) III; cyc-2.1[mkc6(cyc-2.1::3×flag)] IV*, DCL312 *glp-4(bn2) I*, DCL374 *atfs-1(gk3094) V*, DCL419 *gld-1[mkc28(gld-1::mKate2)] I*, DCL430 *gld-1[mkc28(gld-1::mKate2)] I; daf-2(e1370) rsks-1(ok1255) III*, DCL569 *mkcSi13 [sun-1p::rde-1::sun-1 3′UTR + unc-119(+)] II; rde-1(mkc36) V*, XA8205 *aak-2(ok524) X*, XA8222 *daf-2(e1370) rsks-1(ok1255) III*.

Strains in this study were derived from the Bristol N2 background. Nematodes were cultured at 20°C on Nematode Growth Media (NGM) agar plates seeded with *E. coli* OP50 unless otherwise stated (Brenner, 1974).

### Polysomal profiling

Four biological replicates of the wild-type N2 and *daf-2 rsks-1* mutant animals at day 4 of adulthood were collected for polysomal profiling. Around 10,000 worms per sample were homogenized on ice in 350 μl of lysis buffer (300 mM NaCl, 50 mM Tris-HCl [pH 8.0], 10 mM MgCl_2_, 1 mM EGTA, 200 μg / ml heparin, 400 U / ml RNAsin, 1.0 mM phenylmethylsulfonyl fluoride, 0.2 mg / ml cycloheximide, 1% Triton X-100, 0.1% Sodium Deoxycholate) by 60 strokes with a Teflon homogenizer. Each sample was then supplemented with 700 μl lysis buffer and incubated on ice for 30 minutes before centrifuging at 20,000 g for 15 minutes at 4°C. 0.9 ml of the supernatant was applied to the top of a 10-50% sucrose gradient in the high salt resolving buffer (140 mM NaCl, 25 mM Tris-HCl [pH 8.0], 10 mM MgCl_2_) and centrifuged in a Beckman SW41Ti rotor at 38,000 rpm for 90 min at 4°C. Gradients were fractionated using a Teledyne density gradient fractionator with continuous monitoring of absorbance at 252 nm. Translated mRNAs (≥ 2 ribosomes per mRNA) and total RNAs (from the original lysates) were extracted using the Trizol reagent.

### RNA-Seq and bioinformatic analysis

Four biological replicates of translated and total mRNAs from N2 and *daf-2 rsks-1* mutant animals were sent to the University of Minnesota Genomics Center for library construction and pair-ended sequencing with the length of 100 nucleotides, 11.5 million reads per sample on a HiSeq2000 machine (Illumina). Reads were aligned to the *C. elegans* genome (WS220) using the spliced-junction mapper TopHat2 (Kim et al., 2013). Aligned reads were counted per gene using the python script HTseq (Anders et al., 2015). Differential expression and dataset normalization were performed using the Bioconductor package edgeR (Robinson et al., 2010). Normalization in edgeR was adjusted for RNA composition to ensure that highly expressed genes, which consume a large portion of the RNA pool, did not result in the under-sampling of other genes. Dispersion of the gene counts were estimated tag-wise using the Cox-Reid profile-adjusted likelihood method (Cox and Reid, 1987). Only genes with an average counts per million (CPM) of eight or greater across all conditions were considered for differential expression. Differential expression was calculated pairwise between groups using a general linear model and the resulting *p* values were adjusted for multiple testing sing the Benjamin-Hochberg method (Benjamin and Hochberg, 1995). Changes in post-transcriptional processing were identified by comparing the ratio of polysomal-associated mRNA to total mRNA between the wild-type N2 and *daf-2 rsks-1* mutant animals. This is referred to as Differential Polysome Association Ratio (DPAR), with a positive value indicating an increase in the percentage of a particular species of mRNA associated with polysomes in the *daf-2 rsks-1* mutant.

### RNAi by feeding

RNAi experiments were performed by feeding worms *E. coli* strain HT115 (DE3) transformed with either the empty vector L4440 as the control or gene-targeting constructs from the *C. elegans* RNAi Collection (Kamath et al., 2001). Overnight bacterial culture in LB supplemented with Ampicillin (100 μg / ml) at 37°C was seeded onto NGM plates containing IPTG (1mM) and Ampicillin (100 μg / ml) and incubated overnight at room temperature to induce double-stranded RNAs production. Embryos or L1 larvae were placed on RNAi plates and incubated at 20°C until adulthood to score phenotypes.

### Lifespan assay

All lifespan assays were performed at 20°C. To prevent progeny production during lifespan assays, 20 μg / ml (+)-5-fluorodeoxyuridine (FUdR) was added onto NGM plates during the reproductive period (day 1 to 7 of adulthood). The first day of adulthood is Day 1 on survival curves. Animals were scored as alive, dead or lost every other day. Animals that did not respond to gentle touch were scored as dead. Animals that died from causes other than aging, such as sticking to the plate walls, internal hatching or bursting in the vulval region, were scored as lost. Kaplan–Meier survival curves were plotted for each lifespan assay, and statistical analyses (log-rank tests) were performed using the Prism 6 software.

### RNAi screen for lifespan regulators

The primary screen was performed using the strain TJ1060 *spe-9(hc88) I; rrf-3(b26) II*, which showed temperature sensitive sterility, enhanced RNAi sensitivity and normal lifespan. Synchronized TJ1060 L1 larvae were transferred onto plates with RNAi against 112 translationally down-regulated genes at 25°C until animals reached day 1 adulthood for survival assays. For the 39 RNAi treatments that caused larval arrest, synchronized TJ1060 L1 larvae were transferred to the control RNAi plates, and day 1 adult animals were then transferred to those 39 RNAi plates for survival assays. Animals were scored as alive, dead or lost every 4-5 days. RNAi treatments that caused significant lifespan extension (*p* < 0.05, log-rank tests) were used for two rounds of re-tests in the wild-type N2 background at 20°C. RNAi treatments that caused significant lifespan extension in both re-test groups (*p* < 0.05, log-rank tests) were regarded as positive hits.

### Microscopy

*hsp-6p::*GFP transgenic worms were anaesthetized in 50 mM sodium azide and immediately imaged with a Leica MC165 FC dissecting microscope and a Leica DFC450 C digital camera. GLD-1::mKate2 KI animals were anaesthetized in 50 mM sodium azide on agarose pads and imaged with a Zeiss LSM880 confocal microscope.

### Western blot and antibodies

Synchronized day 1 adult animals were collected in the lysis buffer (150 mM NaCl, 1mM EDTA, 0.25% SDS, 1.0% NP-40, 50 mM Tris-HCl [pH7.4], Roche complete protease inhibitors and phosSTOP phosphatase inhibitors) supplemented with the 4 × SDS loading buffer and immediately frozen at −80°C. Samples were boiled for 10 minutes before resolving on precast SDS-PAGE gels (Genscript). Antibodies used in Western blots include monoclonal anti-FLAG (Sigma, 1804), monoclonal anti-Tubulin Alpha (Sigma, T6074), anti-Actin (CST, 4967), anti-Phospho-AMPKα (CST, 2535S) and anti-RPS-0, RPS-3, RPL-5 and RPL-25.2 antibodies (Liu et al., 2018).

### RT-qPCR

Synchronized Day 1 adult animals were collected and frozen in the Trizol reagent (Takara) at −80°C until total RNA extraction using the Direct-zol RNA mini prep kit (ZYMO Research). The cDNA was synthesized by the reverse transcription system (Takara). The SYBR Green dye (Takara) was used for qPCR reactions carried out in triplicates on a Roche LightCycler 480 real-time PCR machine. Relative gene expression levels were calculated using the 2^−∆∆Ct^ method (Livak and Schmittgen, 2001). RT-qPCR experiments were performed at least three times with consistent results using independent RNA preparations. mRNA levels of *pmp-2* were used for normalization.

### Worm tissue micro-dissection

Worms at day 1 adulthood were transferred into the S buffer (100 mM NaCl and 50 mM potassium phosphate [pH 6.0]) on a glass slide. Heads of animals were cut off near the pharynx using syringe needles to collect the intestine and gonad. For RT-qPCR assays, 20-30 gonad or intestine tissues were collected in the Trizol reagent for total RNA extraction. For Western blots, at least 100 gonad or intestine tissues were collected in the protein extraction buffer.

### CRISPR/Cas9 alleles generation

CRISPR engineering to knock-in 3×FLAG at the C-terminal of CYC-2.1 was performed by microinjection using the homologous recombination approach (Dickinson et al., 2013). The injection mix contained two plasmids that drive expression of two different Cas9-sgRNAs (50 ng / μl), a selection marker pCFJ90 (P*myo-2*::mCherry::*unc-54*-3′-UTR) (5 ng / μl, Addgene #19327) and a homologous recombination plasmid (50 ng / μl). To generate the sgRNA plasmids, primers were designed with the CRISPR DESIGN tool (http://crispr.mit.edu) and inserted into the pDD162 vector (Addgene #47549) using the site-directed mutagenesis kit (TOYOBO SMK-101). To generate the homologous recombination plasmid, the 3×FLAG coding sequence was first cloned to replace the GFP coding sequence on vector pPD95.77, and two homologous arms (~1,000 bp each) corresponding to the 5′- and 3′-sides of the insertion site, respectively, were then cloned into the same vector. Successful knock-in events were screened by PCR genotyping from independent F1 transgenic animals’ progenies that did not carry the co-injection markers, and confirmed by Sanger sequencing.

CRISPR engineering with a self-excising drug selection cassette (SEC) was performed to knock-in the mKate2 fluorescent protein to the C-terminal of GLD-1 (Dickinson et al., 2015). The injection mix contained two plasmids that drive expression of two different Cas9-sgRNAs (50 ng / μl), a selection marker pCFJ90 (P*myo-2*::mCherry::*unc-54*-3′-UTR) (5 ng / μl, Addgene #19327) and a repair template FP-SEC vector (50 ng / μl). To generate the sgRNA plasmids, primers were designed with the CRISPR DESIGN tool (http://crispr.mit.edu) and inserted into the pDD162 vector (Addgene #47549) using the site-directed mutagenesis kit (TOYOBO SMK-101). The *ccdB* sequences from the FP-SEC vector pDD287 (Addgene #70685) were replaced with two homologous arms (500-700 bp) to generate the repair template FP-SEC plasmid. Injected animals and their progenies were treated with Hygromycin B (350μg / ml) to select successful knock-in events. Hygromycin B resistant roller animals were tested by PCR genotyping. The SEC was removed by heat shock and homozygous knock-in alleles were confirmed by PCR and DNA sequencing.

### Statistical analysis

Data in bar or scatter graphs were plotted as means ± SD or SEM of at least three independent experiments and evaluated using two-tailed *t* - tests for unpaired samples. Survival curves were evaluated with log-rank tests. *p* < 0.05 was considered significant.

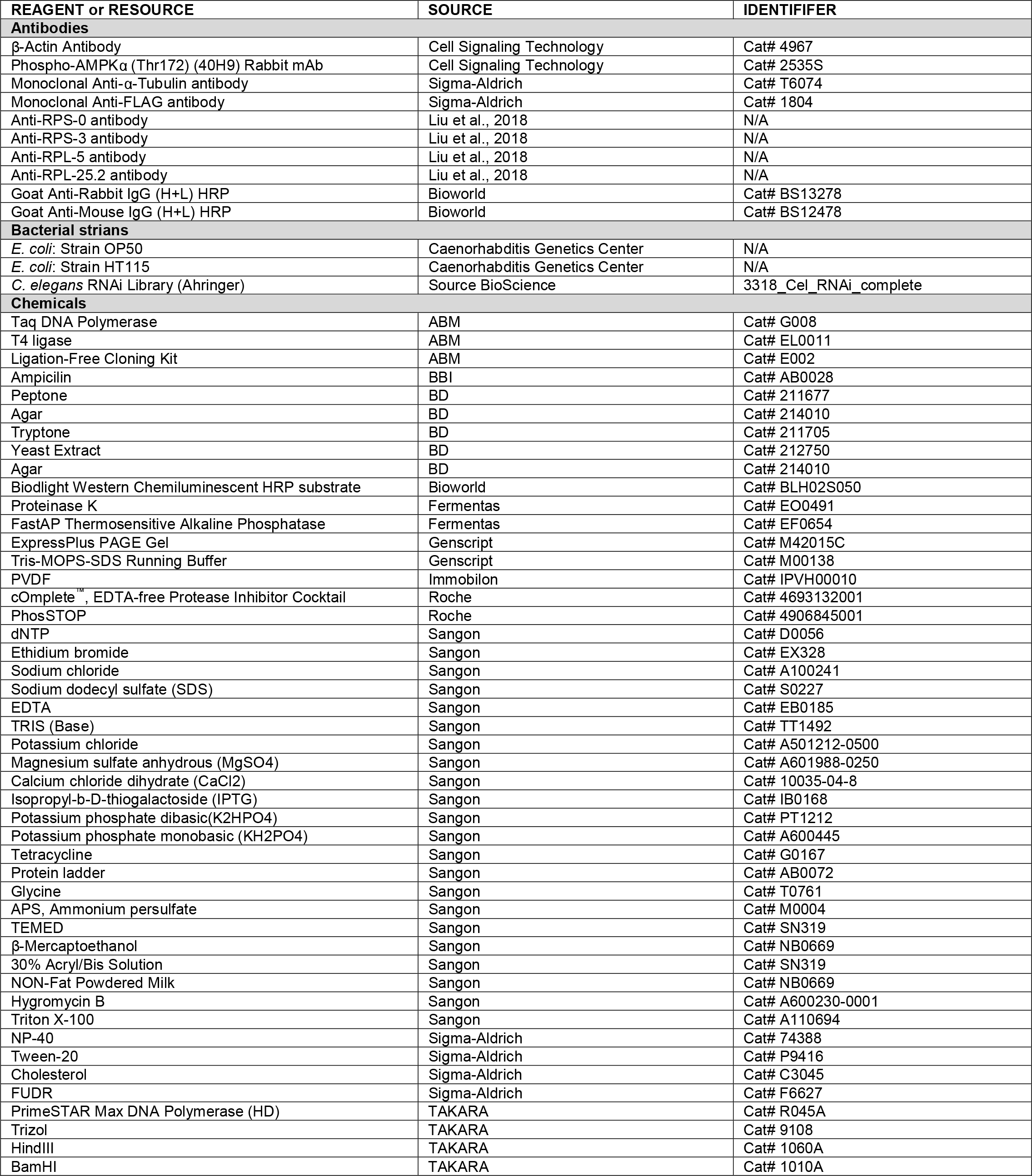

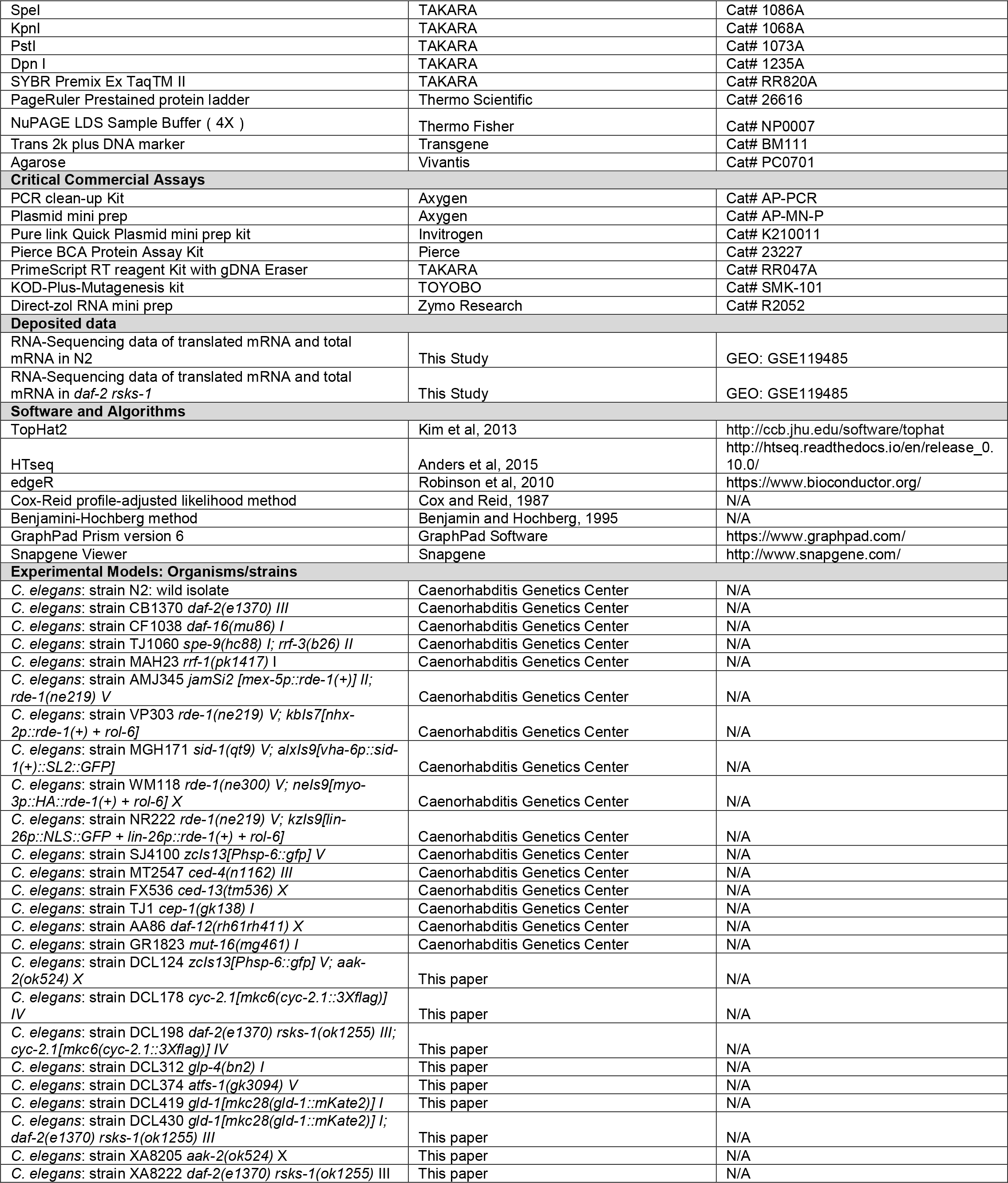

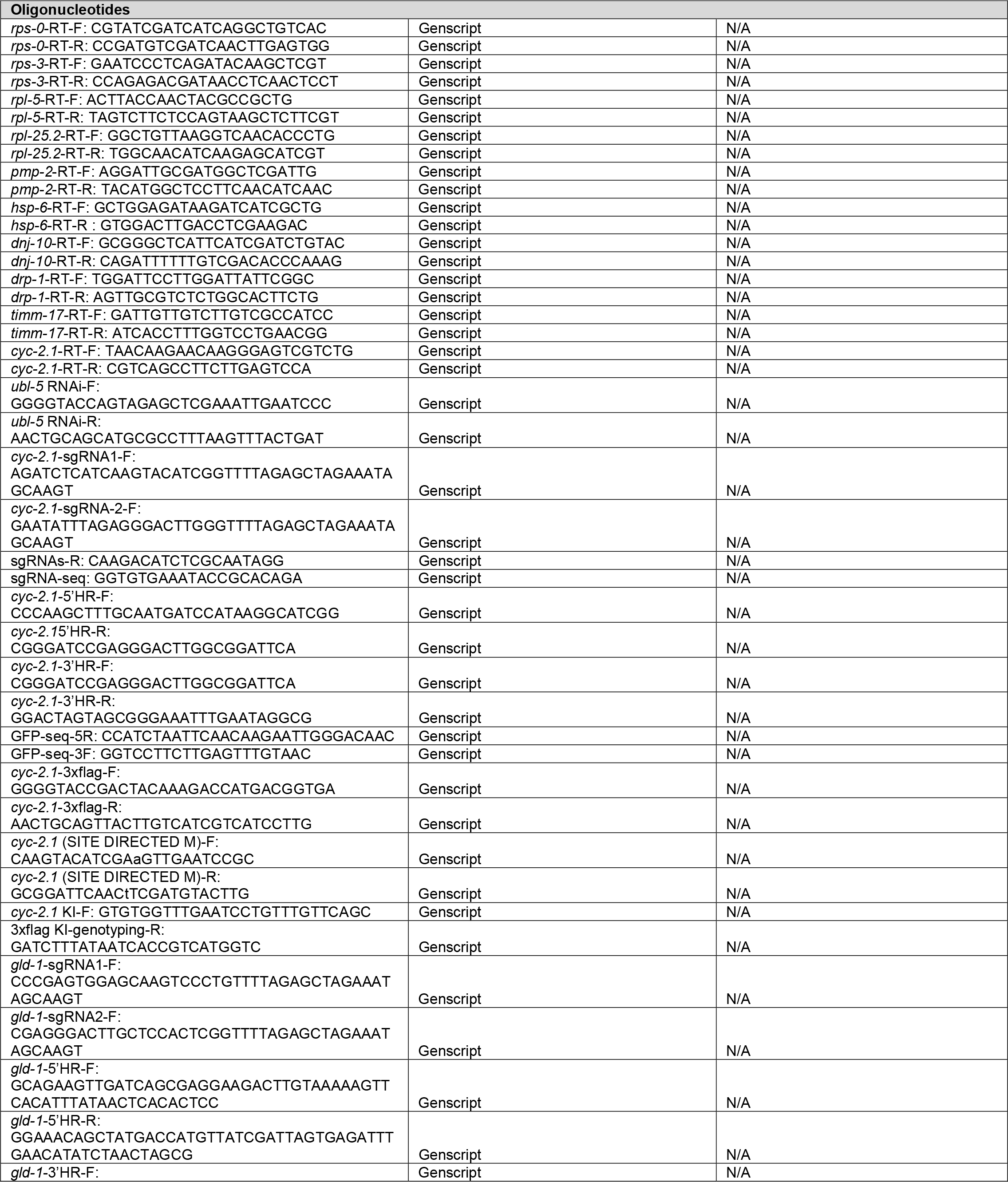

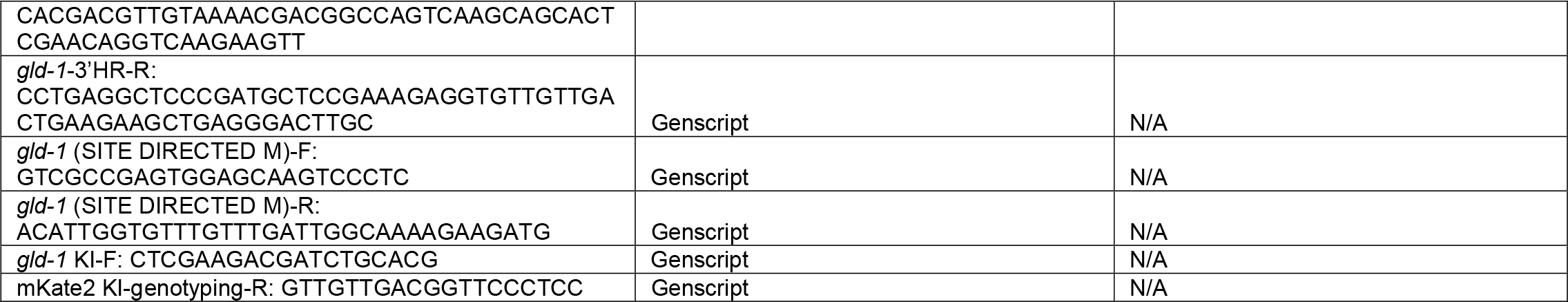

---

**Table S2.** Statistical analyses of lifespan experiments (related to Figure 2, 3, 4, 5)

| Genotype | RNAi | Tissue <sup>a</sup> | Lifespan (days) |  | Percent of control <sup>b</sup> | n <sup>c</sup> | *p* <sup>d</sup> |
| --- | --- | --- | --- | --- | --- | --- | --- |
|  |  |  | Mean | Max |  |  |  |
| *cyc-2.1* RNAi significantly extends lifespan in N2 and *daf-2* mutant (Figure 2A) |  |  |  |  |  |  |  |
| N2 | control |  | 18.9, 20.0 | 29, 29 | / | 72, 88 | / |
| N2 | *cyc-2.1* |  | 32.2, 31.6 | 49, 47 | 170%, 158% | 67, 94 | <0.0001 |
| *daf-2* | control |  | 41.1, 42.2 | 61, 57 | / | 69, 106 | / |
| *daf-2* | *cyc-2.1* |  | 68.8, 68.5 | 105, 97 | 167%, 162% | 82, 156 | <0.0001 |
| *cyc-2.1* RNAi induced lifespan extension requires ATFS-1 and AAK-2 (Figure 2C, F) |  |  |  |  |  |  |  |
| *atfs-1* | control |  | 19.7, 17.6 | 29, 25 | / | 83, 82 | / |
| *atfs-1* | *cyc-2.1* |  | 20.3, 18.6 | 35, 27 | 103%, 106% | 64, 74 | 0.2985, 0.0193 |
| *aak-2* | control |  | 15.9, 16.0 | 21, 21 | / | 71, 55 | / |
| *aak-2* | *cyc-2.1* |  | 15.9, 16.0 | 23, 23 | 100%, 100% | 53, 56 | 0.9647, 0.9295 |
| Effects of tissue-specific *cyc-2.1* RNAi on lifespan (Figure 3A) |  |  |  |  |  |  |  |
| *mkcSi13; rde-1* | control | germline | 17.7, 17.7, 17.5 | 25, 23, 23 | / | 76, 84, 75 | / |
|  | *cyc-2.1* |  | 22.7, 22.0, 22.5 | 35, 31, 33 | 128%, 124%, 129% | 99, 94, 89 | <0.0001, <0.0001, <0.0001 |
| *rde-1; kbls7* | control | intestine | 17.9, 19.4, 16.2 | 25, 27, 23 | / | 62, 94, 51 | / |
|  | *cyc-2.1* |  | 17.9, 17.7, 16.6 | 25, 29, 25 | 100%, 91%, 102% | 74, 98, 78 | 0.9741, 0.0050, 0.4904 |
| *rde-1; kzl9* | control | epidermis | 19.5, 20.6, 20.5 | 29, 29, 27 | / | 114, 101, 70 | / |
|  | *cyc-2.1* |  | 19.0, 19.0, 18.9 | 27, 27, 27 | 97%, 92%, 92% | 99, 112, 105 | 0.1543, 0.0010, 0.0002 |
| *rde-1; nels9* | control | muscle | 19.4, 19.0, 20.7 | 27, 31, 33 | / | 66, 91, 91 | / |
|  | *cyc-2.1* |  | 18.5, 18.8, 21.6 | 29, 29, 33 | 95%, 99%, 105% | 70, 90, 89 | 0.3438, 0.5818, 0.2178 |
| Roles of UPR<sup>mt</sup> pathway transcriptional factors in the regulation of N2 and *daf-2 rsks-1* mutant lifespan (Figure 4D, E, F) |  |  |  |  |  |  |  |
| N2 | control |  | 20.4, 19.8, 19.7 | 29, 27, 25 | / | 64, 86, 58 | / |
|  | *dve-1* |  | 15.8, 14.2, 15.1 | 21, 19, 21 | 78%, 72%, 77% | 83, 97, 93 | <0.0001, <0.0001, <0.0001 |
|  | *ubl-5* |  | 20.6, 19.4, 20.0 | 27, 29, 27 | 101%, 98%, 102% | 77, 95, 92 | 0.8181, 0.6945, 0.1015 |
|  | *atfs-1* |  | 21.3, 20.1, 19.8 | 31, 31, 27 | 104%, 102%, 100% | 73, 93, 49 | 0.2259, 0.2192, 0.7948 |
| *daf-2 rsks-1* | control |  | 55.7, 49.9, 51.7 | 97, 85, 87 | / | 58, 68, 119 | / |
|  | *dve-1* |  | 18.5, 18.7, 22.5 | 31, 27, 41 | 33%, 37%, 43% | 74, 92, 208 | <0.0001, <0.0001, <0.0001 |
|  | *ubl-5* |  | 34.9, 36.2, 32.5 | 59, 55, 55 | 63%, 72%, 63% | 85, 80, 192 | <0.0001, <0.0001, <0.0001 |
|  | *atfs-1* |  | 42.6, 39.5, 44.1 | 69, 59, 83 | 76%, 79%, 85% | 73, 62, 139 | <0.0001, <0.0001, 0.0013 |
| *gld-1* RNAi suppresses super longevity of *daf-2 rsks-1* (Figure 5F) |  |  |  |  |  |  |  |
| *daf-2 rsks-1* | control |  | 65.2, 59.1, 71.3 | 105, 91, 99 | / | 53, 47, 88 | / |
| *daf-2 rsks-1* | *gld-1* |  | 49.6, 52.1, 55.7 | 69, 75, 73 | 76%, 88%, 78% | 43, 45, 75 | <0.0001, 0.0135, <0.0001 |

<sup>a</sup> tissue in which RNAi is effective
<sup>b</sup> changes in mean lifespan compared to the control
<sup>c</sup> numbers of animals scored
<sup>d</sup> p values for log-rank tests

**Figure S1.**
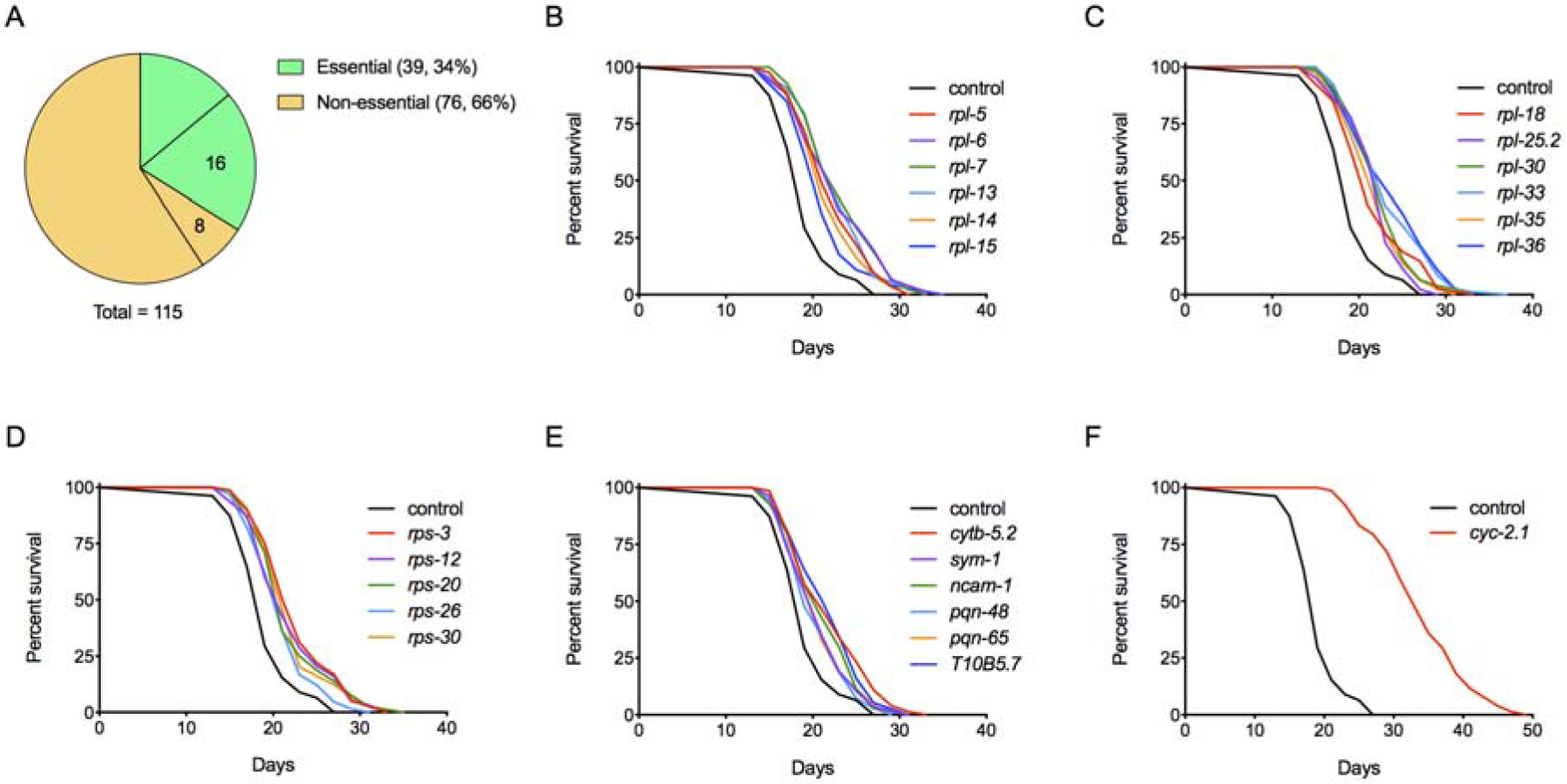
Genes translationally down-regulated in the *daf-2 rsks-1* mutant are enriched with negative regulators of lifespan (related to Table 1) (A) Portions of developmentally essential genes (green) and non-essential genes (yellow) among the 115 genes that are translationally down-regulated in the *daf-2 rsks-1* mutant. Numbers represent the amount of lifespan regulators. (B-D) Survival curves of N2 animals treated with the control RNAi or RNAi against developmentally essential genes that encode various ribosomal subunits. Each RNAi treatment significantly extends lifespan (*p* < 0.05, log-rank tests). (E) Survival curves of N2 animals treated with the control RNAi or RNAi against non-essential genes. Each RNAi treatment significantly extends lifespan (*p* < 0.05, log-rank tests). (F) Survival curves of N2 animals treated with control or *cyc-2.1* by RNAi (*p* < 0.0001, log-rank tests).

**Figure. S2.**
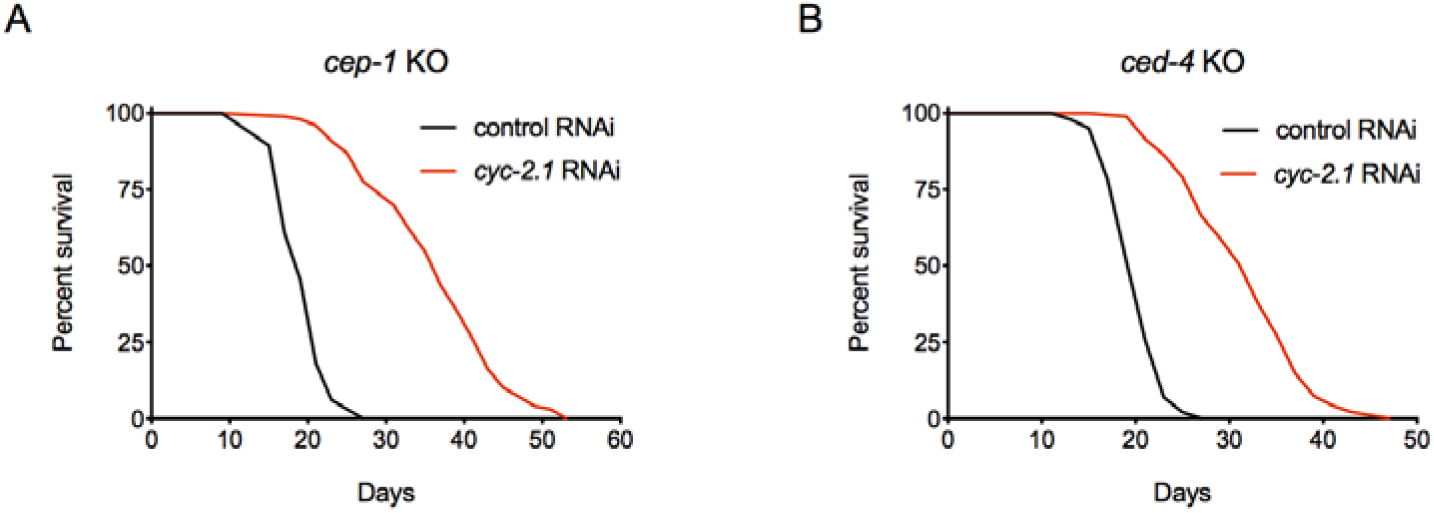
*cyc-2.1* RNAi-induced lifespan extension is independent of CEP-1 and CED-4 (related to Figure 2) (A) Survival curves of the *cep-1* KO mutant treated with either control or *cyc-2.1* RNAi (*p* < 0.0001, log-rank test). (B) Survival curves of the *ced-4* KO mutant treated with either control or *cyc-2.1* RNAi (*p* < 0.0001, log-rank test).

**Figure S3.**
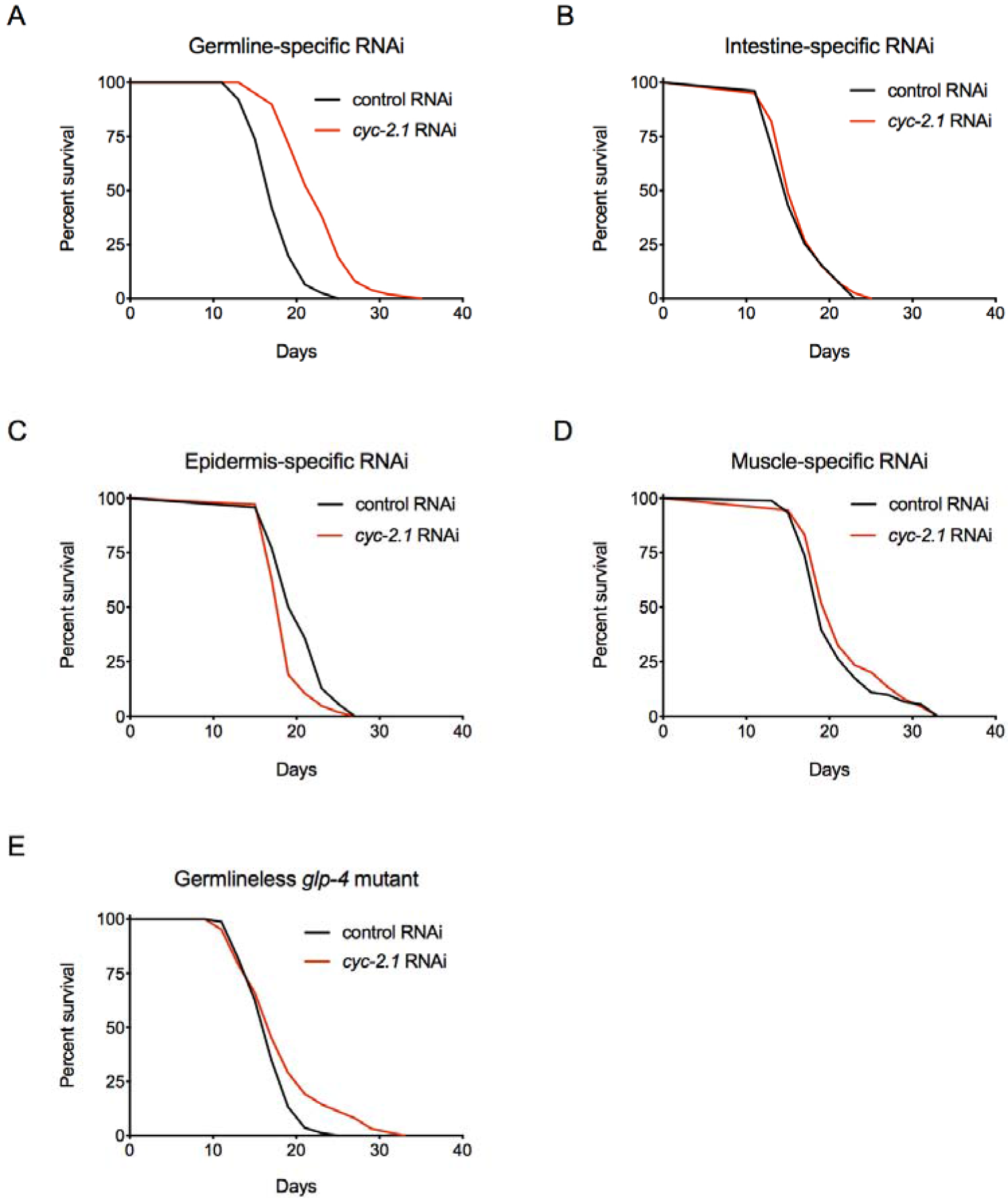
The germline plays an important role in *cyc-2.1* knockdown-induced lifespan extension (related Figure 3) (A-D) Survival curves of animals treated with control RNAi or germline-specific (A), intestine-specific (B), epidermis-specific (C) and muscle-specific (D) *cyc-2.1* RNAi. (E) Survival curves of the germline-less *glp-4* mutant treated with control RNAi or *cyc-2.1* RNAi.

**Figure S4.**
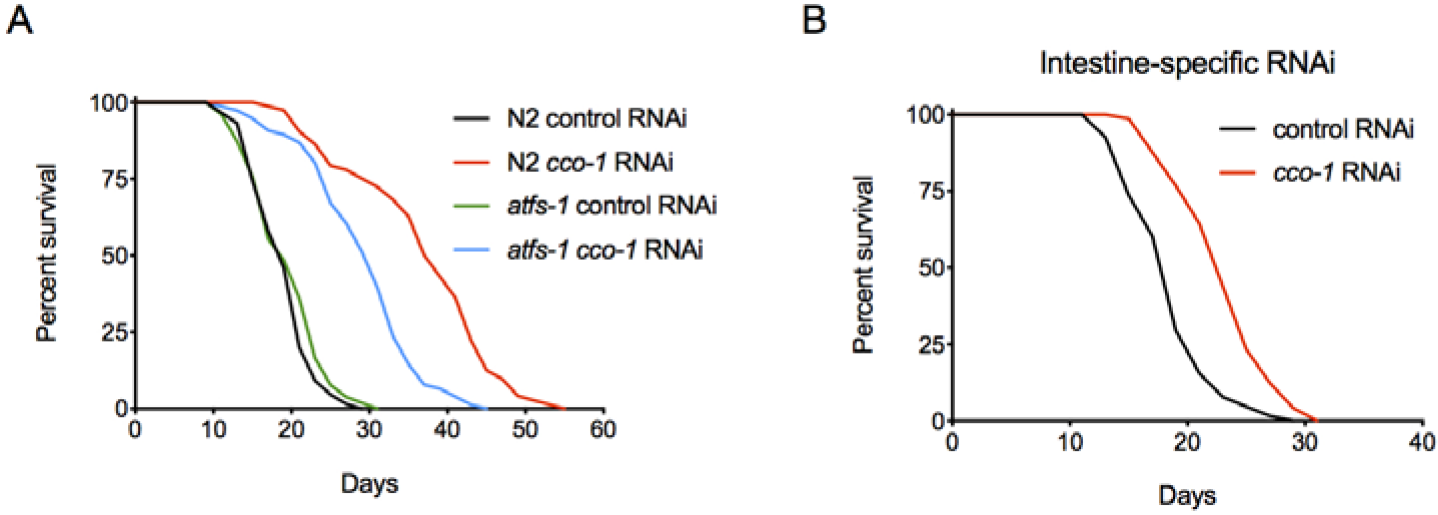
*cco-1* and *cyc-2.1* function through different mechanisms to regulate lifespan (related to Figure 3) (A) Survival curves of N2 and the *atfs-1* KO mutant treated with the control or *cco-1* RNAi. (B) Survival curves of animals treated with the control or intestine-specific *cco-1* RNAi (*p* < 0.0001, log-rank test).

**Figure S5.**
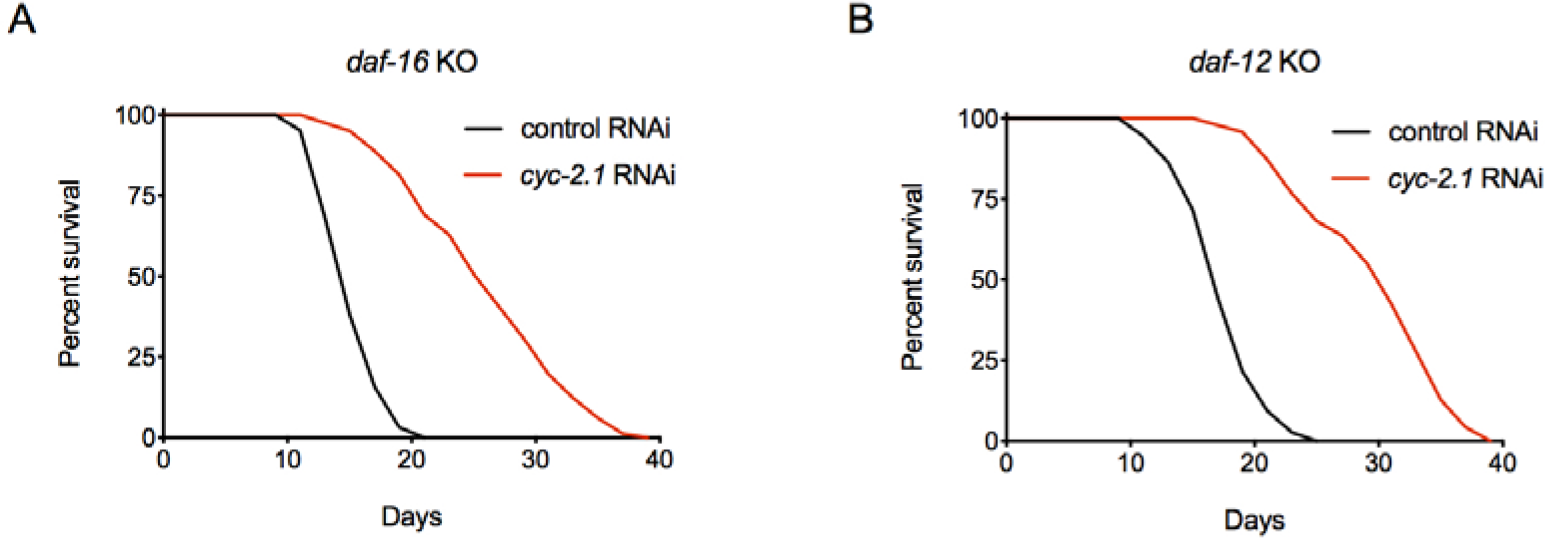
Lifespan extension by *cyc-2.1* RNAi does not require DAF-16 or DAF-12 (related to Figure 3) (A) Survival curves of the *daf-16* KO mutant treated with the control or *cyc-2.1* RNAi. (*p* < 0. 0001, log-rank test). (B) Survival curves of the *daf-12* KO mutant treated with the control or *cyc-2.1* RNAi. (*p* < 0. 0001, log-rank test).

